# The chaperone-client network subordinates cell-cycle entry to growth and stress

**DOI:** 10.1101/485714

**Authors:** David F. Moreno, Eva Parisi, Galal Yahya, Federico Vaggi, Attila Csikász-Nagy, Martí Aldea

## Abstract

The precise coordination of growth and proliferation has a universal prevalence in cell homeostasis. As a prominent property, cell size is modulated by the coordination between these processes in bacterial, yeast and mammalian cells, but the underlying molecular mechanisms are largely unknown. Here we show that multifunctional chaperone systems play a concerted and limiting role in cell-cycle entry, specifically driving nuclear accumulation of the G1 Cdk-cyclin complex. Based on these findings, we establish and test a molecular competition model that recapitulates cell-cycle-entry dependence on growth rate. As key predictions at a single-cell level, we show that availability of the Ydj1 chaperone and nuclear accumulation of the G1 cyclin Cln3 are inversely dependent on growth rate and readily respond to changes in protein synthesis and stress conditions that alter protein folding requirements. Thus, chaperone workload would subordinate Start to the biosynthetic machinery and dynamically adjust proliferation to the growth potential of the cell.

## INTRODUCTION

Under unperturbed conditions of growth cells maintain their size within constant limits, and different pathways have concerted roles in processes leading to growth and proliferation (Cook & Tyers, 2007; Marshall *et al*, 2012; Turner *et al*, 2012). Here we will use the term growth to refer to cell mass or volume increase, while the term proliferation will be restricted to the increase in cell number. Cell growth is dictated by many environmental factors in budding yeast, and the rate at which cells grow has profound effects on their size. High rates of macromolecular synthesis promote growth and increase cell size. Conversely, conditions that reduce cell growth limit macromolecular synthesis and reduce cell size. This behavior is nearly universal and it has been well characterized in bacteria, yeast, diatoms, and mammalian cells of different origins (Aldea *et al*, 2017). A current view sustains that cell cycle and cell growth machineries should be deeply interconnected to ensure cell homeostasis and adaptation, but the causal molecular mechanism is still poorly understood (Lloyd, 2013).

In budding yeast, cyclin Cln3 is the most upstream activator of Start (Tyers *et al*, 1993). Cln3 forms a complex with Cdc28, the cell cycle Cdk in budding yeast, and activates the G1/S regulon with the participation of two other G1 cyclins, Cln1 and Cln2, which contribute to phosphorylate the Whi5 inhibitor thus creating a positive feedback loop that provides Start with robustness and irreversibility (Bertoli *et al*, 2013). The Start network in mammals offers important differences, particularly in the structure and number of transcription factors, but the core of the module is strikingly similar, where Cdk4,6-cyclin D complexes phosphorylate RB and activate E2F-DP transcription factors in a positive feedback loop involving Cdk2-cyclin E (Bertoli *et al*, 2013).

As they are intrinsically unstable, G1 cyclins are thought to transmit growth information for adapting cell size to environmental conditions. The Cln3 cyclin is a dose-dependent activator of Start (Sudbery *et al*,1980; Nash *et al*, 1988; Cross & Blake, 1993) that accumulates in the nucleus due to a constitutive C-terminal NLS (Edgington & Futcher, 2001; Miller & Cross, 2001) and the participation of Hsp70-Hsp40 chaperones, namely Ssa1,2 and Ydj1 (Vergés *et al*, 2007). In addition, Ssa1 and Ydj1 also regulate Cln3 stability (Yaglom *et al*, 1996; Truman *et al*, 2012), and play an essential role in setting the critical size as a function of growth rate (Ferrezuelo *et al*, 2012). In mammalian cells cyclin D1 depends on Hsp70 chaperone activity to form trimeric complexes with Cdk4 and NLS-containing KIP proteins (p21, p27, p57) that drive their nuclear accumulation (Diehl *et al*, 2003).

Molecular chaperones assist nascent proteins in acquiring their native conformation and prevent their aggregation by constraining non-productive interactions. These specialized folding factors also guide protein transport across membranes and modulate protein complex formation by controlling conformational changes (Kampinga & Craig, 2010). Chaperones are involved in key growth-related cellular processes as protein folding and membrane translocation during secretion (Kim *et al*, 2013), and many chaperone-client proteins have crucial roles in the control of growth, cell division, environmental adaptation and development (Gong *et al*, 2009; Taipale *et al*, 2012, 2014). Thus, since chaperones required for Cdk-cyclin activation are also involved in the vast majority of processes underlying cell growth, we hypothesized that competition for shared multifunctional chaperones could subordinate entry into the cell cycle to the biosynthetic machinery of the cell.

Here we show that chaperones play a concerted and limiting role in cell-cycle entry, specifically driving nuclear accumulation of the G1 Cdk-cyclin complex. Ydj1 availability is inversely dependent on growth rate and, based on our findings, we have established a molecular competition model that recapitulates cell-cycle entry dependence on growth rate. As key predictions of the model, we show that nuclear accumulation of the G1 cyclin Cln3 is negatively affected by growth rate in a chaperone-dependent manner, and rapidly responds to conditions that perturb or boost chaperone activity. Thus chaperone availability would act as a G1-Cdk modulator transmitting both intrinsic and extrinsic information to subordinate Start and the critical size to the growth potential of the cell.

## RESULTS

### Nuclear accumulation of the G1 Cdk depends on chaperone activity

Cln3 contains a bipartite nuclear localization signal at its C terminus that is essential for timely entry into the cell cycle (Edgington & Futcher, 2001; Miller & Cross, 2001), and we had found that the Ydj1 chaperone is important for nuclear accumulation of Cln3 (Vergés *et al*, 2007) and for setting the critical size as a function of growth rate (Ferrezuelo *et al*, 2012). Thus, we decided to characterize the role of chaperones as regulators of the G1 Cdk during G1 progression at a single-cell level in time-lapse experiments. Cln3 is too short-lived to be detected as a fluorescent-protein fusion in single cells unless stabilized mutants are used (Liu *et al*, 2015; Schmoller *et al*, 2015). As previously described, mCitrine-Cln3-11A displayed a distinct nuclear signal in most asynchronously growing cells, arguing against the ER retention mechanism that we had proposed previously (Vergés *et al*, 2007). However, this protein is much more stable compared to wild-type Cln3, and transient retention at the ER would likely be obscured by accumulation of abnormally stable mCitrine-Cln3-11A in the nucleus. Thus, we decided to test whether nuclear accumulation of this stabilized protein was still dependent on Ydj1 by carefully measuring nuclear and cytoplasmic fluorescence levels (Fig EV1). While overall levels as determined by immunoblotting were not altered, nuclear mCitrine-Cln3-11A levels strongly decreased in Ydj1-deficient cells (Fig 1A-C), which confirmed previous observations obtained with 3HA-tagged wild-type Cln3 by immunofluorescence (Vergés *et al*, 2007). Moreover, while nuclear levels of Cdc28-GFP in late G1 cells were also negatively affected by deletion of *YDJ1*, overexpression of both Ydj1 and Ssa1 significantly increased the nuclear to cytoplasmic ratio of Cdc28-GFP both in mid- and late-G1 cells (Fig 1D). Likely due to the fact that Cdc28 is present at much higher levels than Cln3 (Tyers *et al*, 1993; Cross *et al*, 2002) differences in the steady-state nuclear levels of Cdc28 when comparing wild-type and Ydj1-deficient or overexpressing cells were only modest (Fig 1D). On the other hand, we cannot discard the effects of Cln1,2 cyclins synthesized at Start by the transcriptional feedback loop. Thus, we decided to analyze directly the import kinetics of Cdc28-GFP in G1 cells by nuclear FLIP (Figs 1E,F and EV2). We found that, while being extremely dependent on Cln3 (Fig 1G), the nuclear import rate of Cdc28-GFP decreased in Ydj1-deficient cells and was clearly impaired when chaperone function was compromised by azetidine 2-carboxylic acid (AZC), a proline analog that interferes with proper protein folding (Trotter *et al*, 2001). By contrast, a 2NLS-GFP control did not show significant differences in nuclear localization or import kinetics in cells lacking Ydj1, Cln3 or in the presence of AZC (Fig 1C and H). These data are consistent with a chaperone-dependent mechanism that drives nuclear import of the Cdc28-Cln3 complex in G1 for the timely execution of Start.

**Figure 1.**
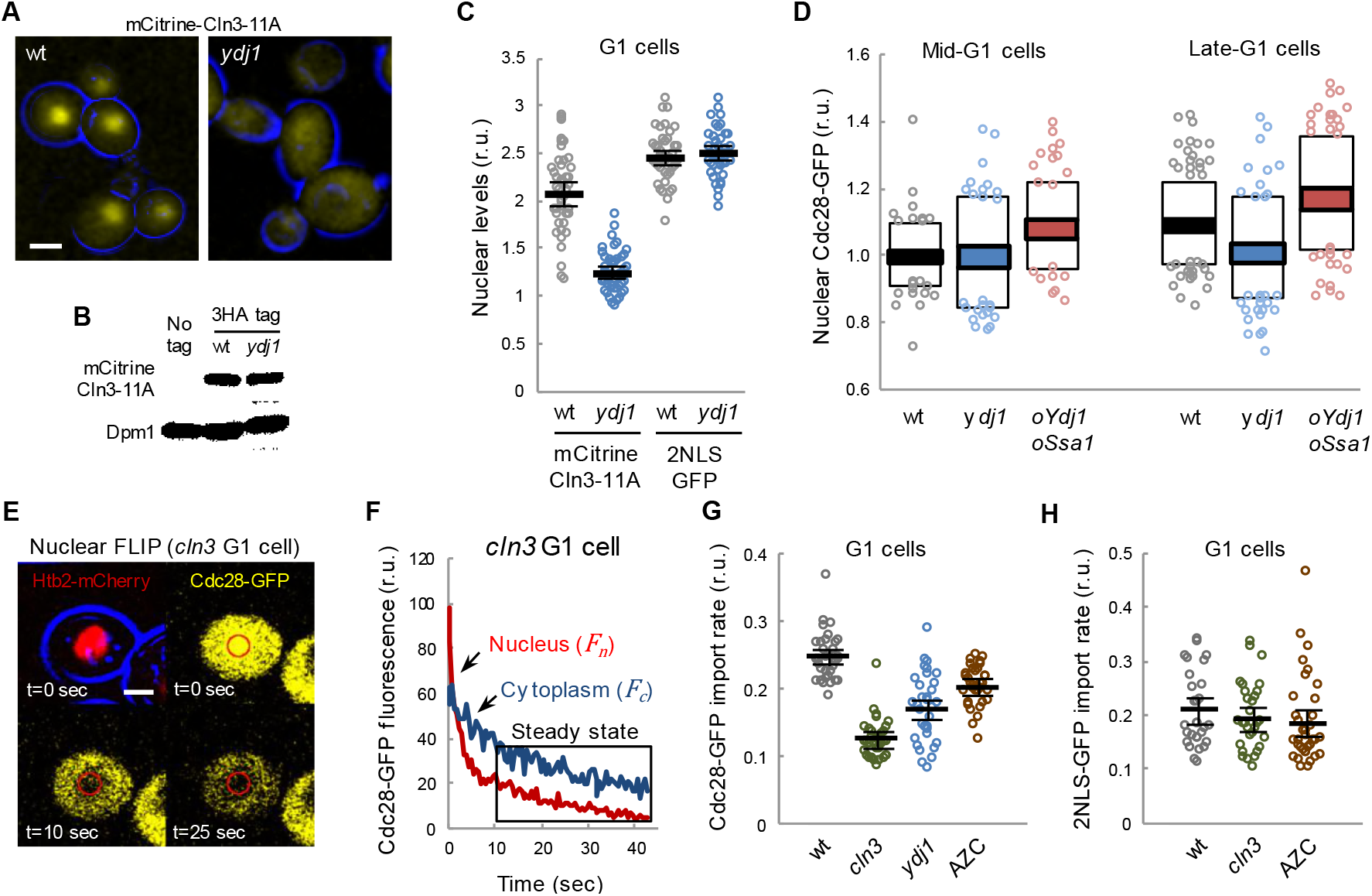
Nuclear accumulation of the G1 Cdk depends on chaperone function. A Images corresponding to wild-type (wt) and Ydj1-deficient (*ydj1*) cells expressing mCitrine-Cln3-11A. Bar is 2 μm. B Immunoblotting analysis of 3HA-tagged mCitrine-Cln3-11A levels in wild-type and *ydj1* cells. Dpm1 is shown as loading control. C Nuclear to cytoplasmic mCitrine-Cln3-11A and 2NLS-GFP ratios for individual wild-type and *ydj1* G1 cells. Mean (N=30, thick lines) and confidence limits (α=0.05, thin lines) for the mean are also plotted. D Cdc28-GFP wild-type (wt), Ydj1-deficient (*ydj1*) or overexpressing Ydj1 and Ssa1 (*oYdj1 oSsa1*) cells were analyzed by time-lapse microscopy during G1 progression. Mean (N>100) nuclear to cytoplasmic Cdc28-GFP ratios are plotted with respective standard deviation (white boxes) and confidence (α=0.05, colored boxes) intervals for the mean at either mid (36-45 min before Start) or late G1 (18-21 min before Start). E Analysis of Cdc-28-GFP import by nuclear FLIP. A representative *cln3* cell expressing Cdc28-GFP and Htb2-mCherry at different times during nuclear photobleaching is shown. Bar is 1 μm. F A representative nuclear FLIP output of Cdc28-GFP in a Cln3-deficient G1 cell showing fluorescence decay in nuclear and cytoplasmic compartments. G Cdc28-GFP import rates in wild-type (wt), Cln3-deficient (*cln3*), and Ydj1-deficient (*ydj1*) single cells in G1 phase. Wild-type cells treated with AZC are also shown. Mean values (N>30, thick lines) with confidence limits for the mean (α=0.05, thin lines) are also plotted. H 2NLS-GFP-EBD import rates in wild-type (wt) and Cln3-deficient (*cln3*) G1 single cells after in the presence of 1μM auxin. Wild-type cells treated with AZC are also shown. Mean values (N>30, thick lines) with confidence limits for the mean (α=0.05, thin lines) are also plotted.

### Multifunctional chaperones have a limiting role in setting cell size at budding

If chaperones have a role in coordinating cell growth and Start machineries, chaperone availability ought to be limiting for cell-cycle entry. Thus, we decided to test this proposition by increasing the gene dosage of different chaperone sets in low-copy centromeric vectors. In addition to the Hsp70 system (Ssa1 and Ydj1), we analyzed the effects of key components of the Hsp90 system (Hsc82 and Cdc37), which is important for holding the Cdk in a productive conformation for binding cyclins (Vaughan *et al*, 2006), and the segregase Cdc48 complex (Cdc48, Ufd1, and Npl4), which prevents degradation of ubiquitinated Cln3, and concurrently stimulates its ER release and nuclear accumulation to trigger Start (Parisi *et al*, 2018). Gene duplication in centromeric vectors increased chaperone gene expression by 1.5-to 2-fold in a specific manner (Fig EV3A). Notably, although expression levels increased only modestly, additional copies of chaperone genes caused an additive reduction of the budding volume in asynchronous cultures (Fig 2A) and newborn daughter cells (Fig 2B). This effect was barely observed in large cells deficient for Cln3, and was totally abolished in cells lacking Cln3 as well as Whi5 and Stb1, two key inhibitors of the Start network (Fig 2B). This triple mutant displays unaltered average budding volume compared to wild type (Wang *et al*, 2009), but has lost most of the dependency on growth rate (Ferrezuelo *et al*, 2012). In contrast, the effect was still clear in cells lacking Whi7, an inhibitor of Start that acts at an upstream step (Yahya *et al*, 2014). Finally, we observed a comparable reduction of the budding volume when genes of the three chaperone systems were duplicated together in an artificial chromosome (Fig 2C). Doubling times of cells with vectors expressing the different chaperone sets were not significantly different from wild-type cells. Also, we had found no differences in the growth rate during G1 when comparing wild-type and Ydj1/Ssa1 overexpressing cells (Ferrezuelo *et al*, 2012), which rules out indirect effects on cell size by growth rate changes. In addition, protein levels and phosphorylation status of Cln3 were not affected by chaperone gene dosage (Fig EV3B), pointing to a limiting role of these chaperone sets in Cdk-cyclin complex activity and/or localization.

**Figure 2.**
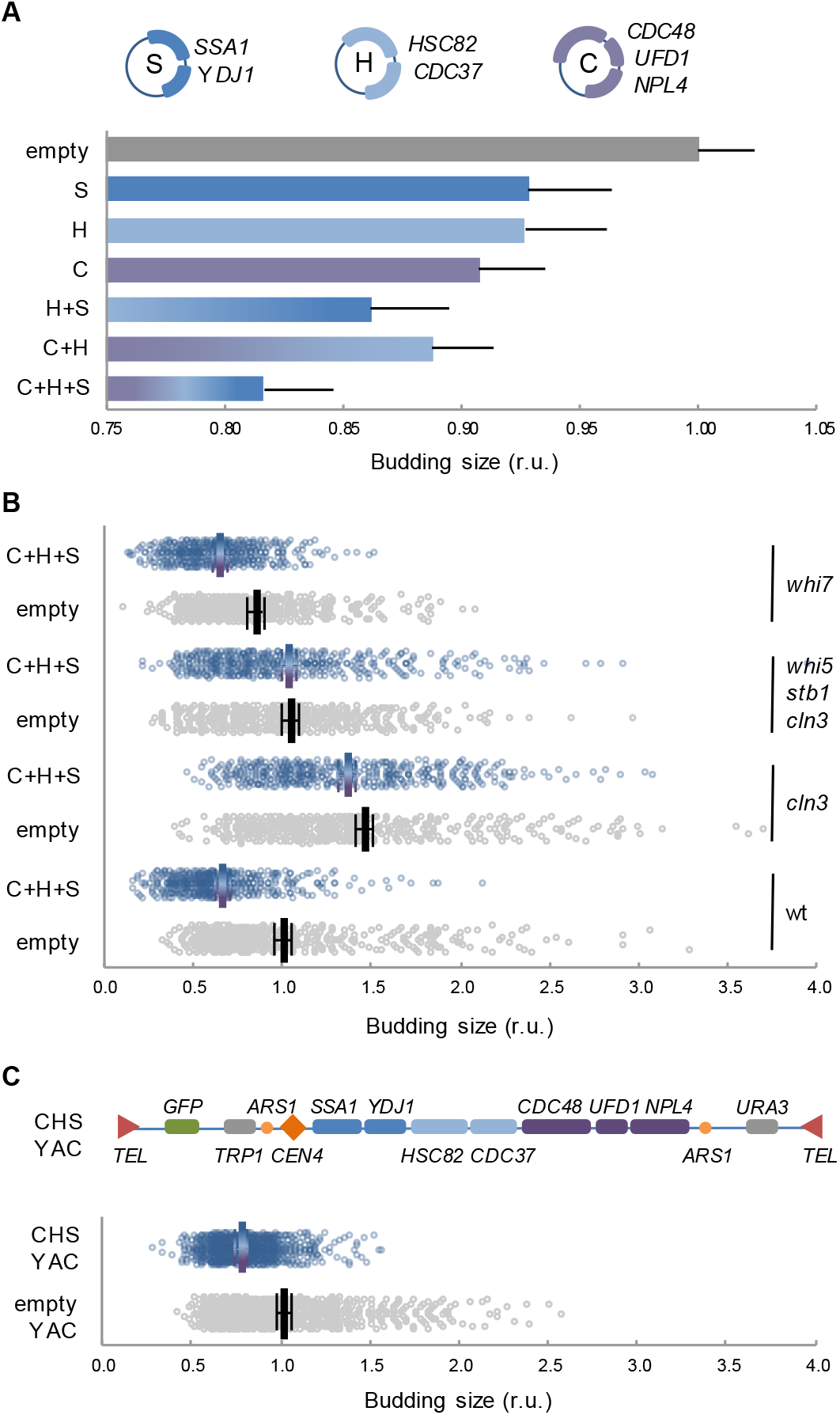
Chaperones of the Hsp70, Hsp90 and Cdc48 systems are limiting for cell cycle entry. A Budding volume of yeast cells transformed with the indicated combinations of compatible centromeric vectors encoding chaperones of the Hsp70 (S, *SSA1* and *YDJ1*), Hsp90 (H, *HSC82* and *CDC37*), and Cdc48 (C, *CDC48, UFD1* and *NPL4*) systems. Individual budding volumes were determined and made relative to the mean value for wild-type cells transformed with the corresponding empty vectors. Mean values (N>200) and confidence limits (α=0.05, thin vertical lines) for the mean are plotted. B Yeast cells with indicated genotypes were transformed with empty or compatible centromeric vectors encoding the three chaperone systems (C+H+S) and budding volumes were determined as in panel A. Individual data (N>400), mean values (thick vertical lines) and confidence limits (α=0.05, thin vertical lines) for the mean areplotted. C Budding volume of yeast cells transformed with an artificial chromosome encoding the three chaperone systems (CHS YAC) or empty vector. Budding volumes were determined as in panel A. Individual data (N<400), mean values (thick vertical lines) and confidence limits (α=0.05, thin vertical lines) for the mean are plotted.

### Molecular competition for chaperones predicts cell size dependency on growth rate

As they are associated with client proteins mostly in a transient manner, the level of free chaperones might be inversely dependent on protein synthesis and trafficking rates, thus constituting a simple mechanism to report growth kinetics to the Start network and, hence, modulate cell size as a function of growth rate. To test this notion, we developed a mathematical model (Fig 3A) wherein protein synthesis and G1 Cdk-cyclin complex assembly compete for limiting amounts of Ydj1, the best characterized chaperone in terms of regulating Cln3. Ydj1 plays an activating role by releasing Cln3 from the ER during G1 (Vergés *et al*, 2007), but is also important for efficient degradation of Cln3 by Cdc28-dependent and autoactivated phosphorylation (Yaglom *et al*, 1996, 1995). Taken together, these data point to the notion that Ssa1/Ydj1 chaperones contribute to both proper Cln3 folding (i.e. binding Cdc28 in a productive conformation) and its release from the ER where the segregase Cdc48 plays also a key role (Parisi *et al*,2018). Since these regulatory steps are likely related at the molecular level, we opted for treating them as a single event in the competition model. On another point, although the specific affinity of Ydj1 for the various Cln3 domains was similar to other proteins (Fig EV4), the number of client proteins being synthesized at any given time is in overwhelming excess relative to those of Ydj1 (Gong *et al*, 2009) and especially of Cln3, which is present at very low levels (Tyers *et al*, 1993; Cross *et al*, 2002).

**Figure 3.**
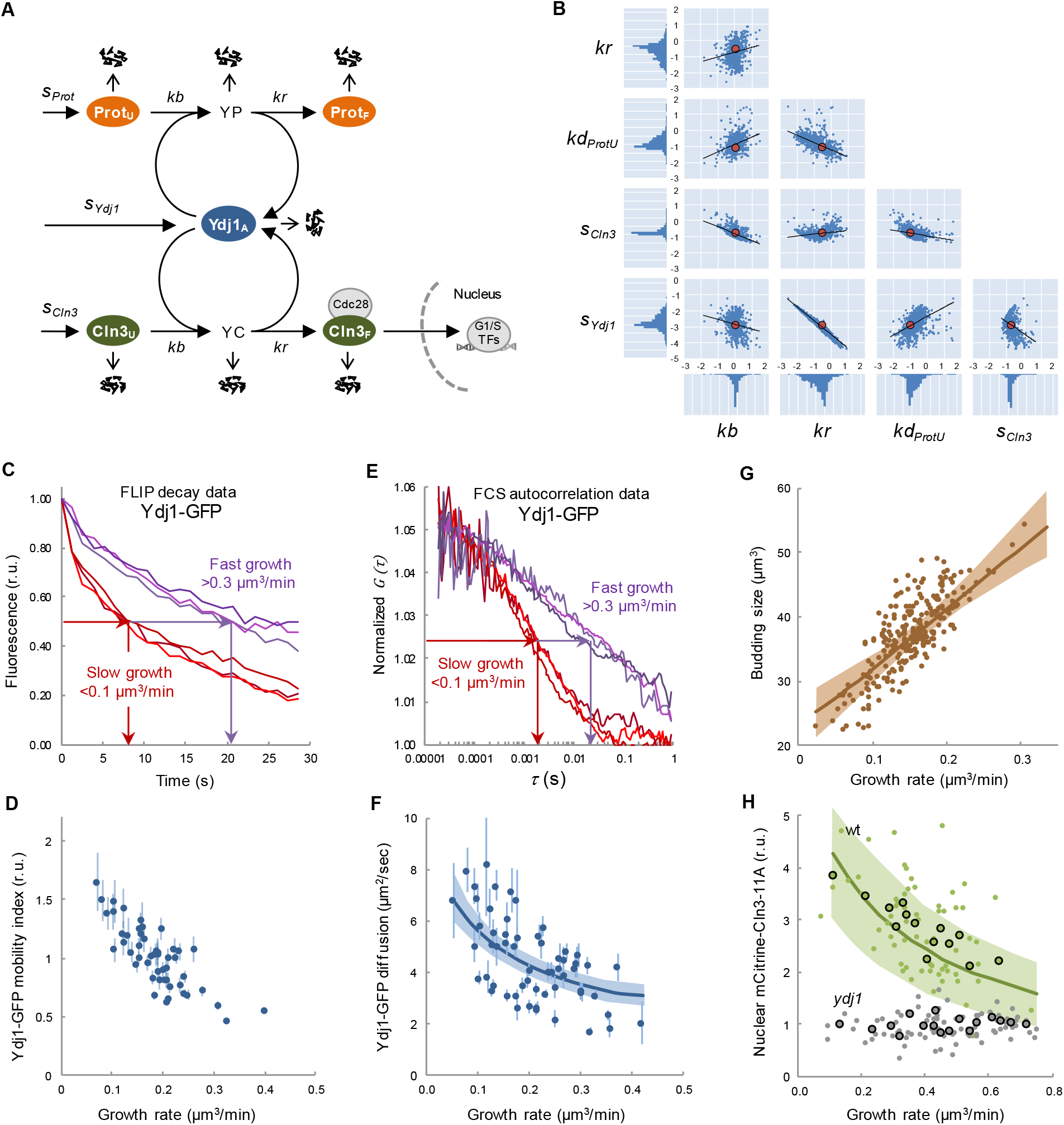
The molecular competition model, chaperone availability and nuclear accumulation of Cln3. A Scheme of the chaperone competition model connecting protein synthesis and Cln3 folding and complex formation with Cdc28. Chaperones associate to the unfolded protein (ProtU) mostly in a transient manner until the properly folded protein (Prot_F_) is released. Thus, protein synthesis rate would be a key determinant of the level of available Ydj1 (Ydj1_A_) in the free pool and, as it is essential in proper folding and release of Cln3 (Cln3_F_), it would in turn govern the rate of Whi5 phosphorylation in the nucleus for triggering Start. Variables and parameters used in the model are indicated. All key components of the competition model are subject to specific rates of degradation (open arrows) as described in Materials and Methods. B Distribution of parameters of the model fitted to experimental data of Ydj1 diffusion and critical volume as a function of growth rate. Parameter sets sampled using Markov Chain Monte Carlo (small blue circles) are plotted in log10 space as well as the corresponding mode values (large red circles) and regression lines. One-dimensional histograms of parameter values are also plotted adjacent to the axes. C Yeast cells expressing Ydj1-GFP were analyzed by FLIP in time-lapse experiments to determine also growth rate at a single-cell level. Individual FLIP decay data are plotted for three fast (>0.3 μm3/min, purple lines) and three slow (<0.1 μm3/min, red lines) growing cells. D Ydj1-GFP mobility indexes (circles) were obtained from FLIP decay curves as shown in panel C and plotted as a function of growth rate with the corresponding confidence limits (α=0.05). E Yeast cells expressing Ydj1-GFP were analyzed by FCS in time-lapse experiments to determine also growth rate at a single-cell level. Individual FCS autocorrelation data are plotted for three fast (>0.3 μm3/min, purple lines) and three slow (<0.1 μm3/min, red lines) growing cells. F Ydj1-GFP diffusion coefficients (circles) were obtained from FCS autocorrelation functions as shown in panel E and plotted as a function of growth rate with the corresponding standard error. The mean fit produced by the full ensemble of parameter sets shown in panel B is plotted as a line with one standard deviation intervals. G The chaperone competition model predicts the critical size being a function of growth rate at the single-cell level. Experimental budding volumes as a function of growth rate (closed circles) and the mean fit as in panel F are shown. H Nuclear to cytoplasmic ratios for mCitrine-Cln3-11A from G1 wild-type (wt, green, N=68) and Ydj1-deficient (*ydj1*, gray, N=85) cells as a function of growth rate. Mean values (large circles) and confidence limits (α=0.05) for binned (5 cells/bin) data are also shown. A simulation of Cln3_F_ was obtained with parameter set 3114 within a 4-fold range of *kd* and *kr*, and the resulting mean (green line) is plotted with one standard deviation intervals.

In our model, the level of available, unbound Ydj1 (Ydj1_A_) is a key variable and, since client-engaged chaperones offer a reduced mobility (Lajoie *et al*, 2012; Saarikangas *et al*, 2017), we decided to use FLIP and FCS to obtain a mobility index of fully functional Ydj1 and Ssa1 fusions to GFP as reporter of their availability in single cells. First we tested the validity of this methodological approach by compromising chaperone function with AZC, which induces misfolded protein accumulation with chaperones into disperse cellular aggregates (Escusa-Toret *et al*, 2013), thus reducing levels of soluble available chaperones. While GFP mobility remained unaffected, both Ssa1 and Ydj1 fusions to GFP decreased their mobility upon AZC treatment (Fig EV5).

Individual genetically-identical cells display a large variability in multifactorial processes such as gene expression and growth (Blake *et al*, 2003; Ferrezuelo *et al*, 2012) and we reasoned that, if the molecular competition model is correct, endogenous variability in growth rate should have an effect on chaperone availability at the single-cell level. We found that Ydj1-GFP diffusion was correlated negatively with growth rate in single G1 cells by both FLIP (Fig 3C and D) and FCS (Fig 3E and F) analysis, and at a population level in media sustaining different growth rates (Fig EV6A). The chaperone-competition model perfectly fitted the dependence of Ydj1 availability on growth rate (Fig 3F) and, more important, it also simulated very closely the increase of the budding volume with growth rate (Ferrezuelo *et al*, 2012) (Fig 3G). As expected from a competition framework, the fitted parameter sets produced negative correlations between available Ydj1 and unfolded target proteins (Fig EV6B), and a positive correlation between the level of Ydj1 in complexes with either Cln3 or all other proteins (Fig EV6C). Interestingly, the competition model produced acceptable fits with parameters spanning several orders of magnitude (Fig 3B), which underlines the robustness of the chaperone competition design in predicting growth-rate dependent chaperone availability and cell size, letting parameters to be adapted for subjugating Start to cellular processes other than growth. In summary, our experimental results and modeling simulations support the notion that growth rate modulates levels of available chaperones.

### Growth rate and protein synthesis modulate accumulation of Cln3 in the nucleus

The molecular competition model predicted that increasing growth rate would decrease available levels of free chaperones (Ydj1_A_) and, hence, decrease the steady-state level of folded free Cln3 (Cln3_F_) (Fig EV6D and E). More precisely, the fraction of Ydj1 bound to client proteins (YP+YC) increases at higher growth rates. As a consequence, the fraction of available free Ydj1 (Ydj1_A_) drops and the rate at which proteins can be folded is reduced, which in turn increases the fraction of unfolded proteins (ProtU and Cln3_U_). Due to its intrinsic instability, folded Cln3 (Cln3_F_) levels are extremely more sensitive to the effects of Ydj1 compared to all other proteins. Indeed, whereas the total concentration of mCitrine-Cln3-11A in G1 remained unaltered on average at different growth rates (Fig EV6G), nuclear levels of mCitrine-Cln3-11A displayed a significant negative correlation (p=2.10^-4^) with growth rate very similar to that predicted by the model (Fig 3H). Notably, this negative correlation was totally lost in Ydj1-deficient cells (Fig 3H). In summary, higher growth rates reduce the nuclear levels of Cln3 at a given volume in a chaperone-dependent manner.

Next, with the purpose of simulating lower growth rates, we used cycloheximide (CHX) to decrease protein synthesis and relieve chaperones temporarily from the load of nascent polypeptides (Fig 4A and B). As expected, CHX inhibited incorporation of puromycin in cell-free extracts (Fig 4C) but it increased the rate at which heat-treated luciferase was renatured and became active, indicating that both protein refolding and synthesis compete for chaperones in cell extracts. Next, we tested the effects of CHX at 0.2 μg/ml, a sublethal concentration that does not activate the environmental stress response (Jacquet, 2003; Trotter *et al*, 2002), and found that the protein synthesis rate displayed a 5.9-fold decrease in live cells (Fig EV7). In agreement with our model prediction (Fig 4B) this sublethal dose of CHX increased the average diffusion coefficient of Ydj1-GFP as measured by FCS (Fig 4D) and FLIP (Fig 4E). Temporary perturbations of chaperone availability should in turn have an effect in the nuclear accumulation of Cln3 (Fig 4B). In order to quantify endogenously expressed Cln3-3HA in immunofluorescence images we used a semiautomated method that analyzes both cytoplasmic and nuclear compartments in fixed yeast cells (Yahya *et al*, 2014). Notably, we found that relative nuclear levels of Cln3-3HA rapidly increased after addition of CHX (Fig 4F and G), decreasing at later times as predicted by the model (Fig 4B), and this transient increase fully depended on Ydj1. However, CHX has been shown to increase cell size at Start (Jorgensen *et al*, 2004). This apparent discrepancy could originate from the different short- and long-term effects of CHX, i.e. increasing Ydj1 mobility and nuclear localization of Cln3 at very short times (less than one minute), but eventually decreasing G1 cyclin levels, which is what would finally result in a larger cell size. In the overall, these data support the notion that chaperone availability transmits growth and protein synthesis rate information to modulate the rate at which the G1 Cdk-cyclin complex is properly formed and accumulates in the nucleus.

**Figure 4.**
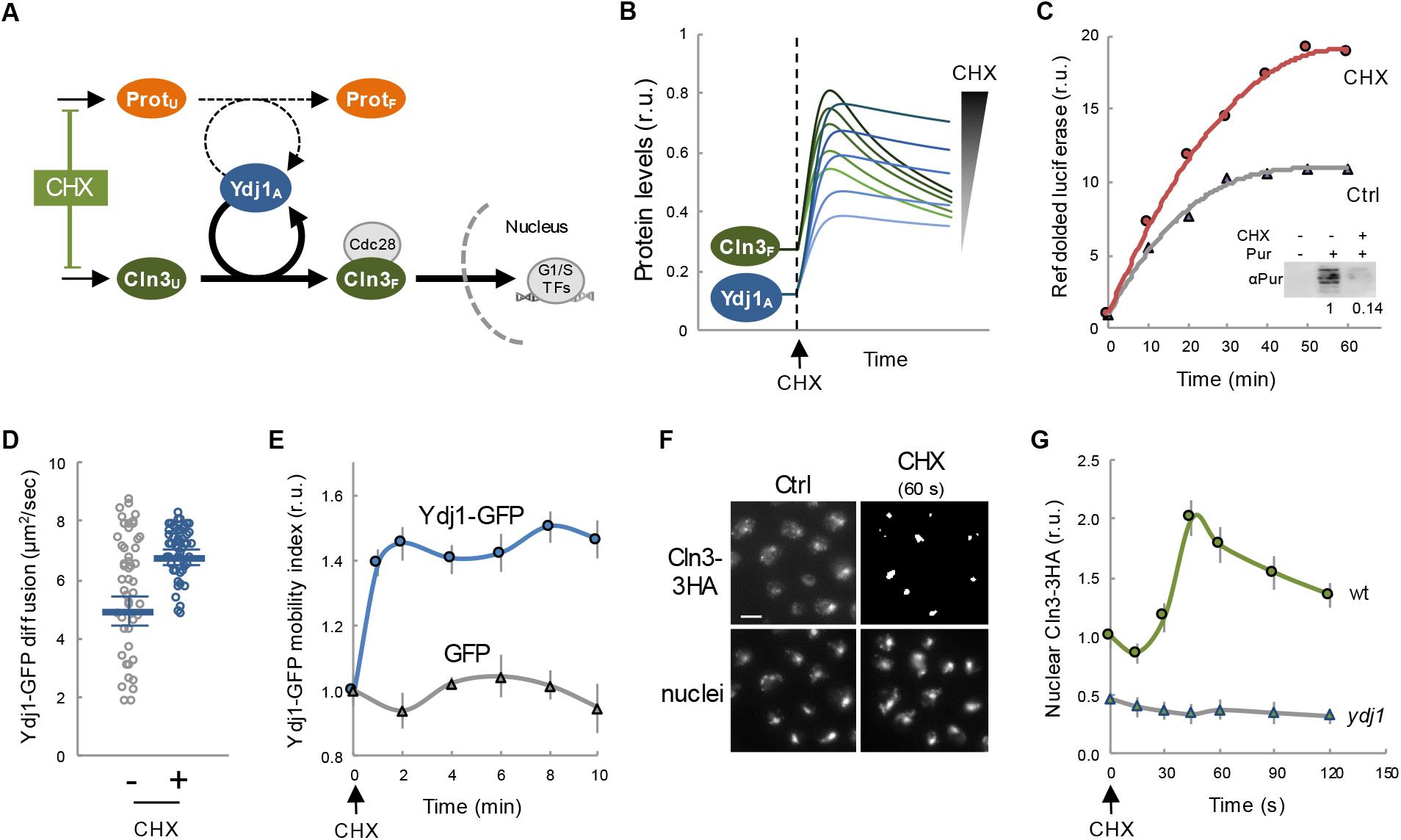
Protein synthesis is a key determinant of chaperone availability and nuclear accumulation of Cln3. A The competition model predicts that reducing the protein synthesis rate would decrease the requirements of Ydj1 chaperone in folding of new proteins (Prot_F_), and increase the level of available chaperone (Ydj1_A_) and free Cln3 (Cln3_F_). B Prediction of available chaperone (Ydj1_A_) and free Cln3 (Cln3_F_) levels as a function of time after CHX addition. Simulations were produced by using parameter set 3114 and varying (3, 4.5, 6, 9, or 12-fold) reductions in protein synthesis rates around the experimental value (Fig EV7). C Refolded luciferase activity as a function of time in yeast cell extracts treated or not with 20 μg/ml CHX in the presence of an ATP-regenerating system. Inset: Immunoblot analysis of puromycin incorporation in cell-free extracts used for refolding analysis in the absence or presence of 20 μg/ml CHX. Numbers refer to relative incorporation levels as measured by densitometric analysis. D Yeast cells expressing Ydj1-GFP were analyzed by FCS before (-) or 5 to 10 min after adding a sublethal dose of CHX at 0.2 μg/ml (+). Individual protein diffusion coefficients are plotted (N>50). Mean values (thick lines) and confidence limits for the mean (α=0.05, thin lines) are also shown. E Ydj1-GFP and GFP mobility assayed by FLIP at the indicated time points after CHX addition at 0.2 μg/ml. Relative mean values and confidence limits (α=0.05) for the mean are shown. F Nuclear levels of Cln3-3HA by immunofluorescence before or 60 sec after addition of CHX at 0.2 μg/ml. G Nuclear accumulation of Cln3-3HA in asynchronous individual wild type (wt) and Ydj1-deficient (*ydj1*) yeast cells before or at the indicated times after CHX addition as in panel F. Relative mean values (N>200) and confidence limits (α=0.05, thin horizontal lines) for the mean are shown.

### Stress conditions that decrease chaperone availability prevent accumulation of Cln3 in the nucleus

Yeast cells respond to stress conditions as diverse as high temperature, high osmolarity or abnormal levels of unfolded proteins in the ER, by highly conserved transcriptional programs that increase chaperone expression to protect damaged proteins from aggregation, unfold aggregated proteins, and refold damaged proteins or target them for efficient degradation (De Nadal *et al*, 2011). Thus, stresses are assumed to cause a temporary deficit in chaperone availability. Hsf1, the key transcriptional activator of the heat shock response, is inhibited by chaperones of the Hsp70 and Hsp90 systems, and it has been proposed that the accumulation of unfolded or damaged proteins would readily titrate the chaperone machinery from Hsf1, allowing derepression of the transcription factor (Verghese *et al*, 2012). Supporting this notion, Ydj1-assisted Ssa1 chaperone is targeted to and accumulates in protein aggregates (Mogk *et al*, 2018) after heat stress. A similar titration mechanism has been proposed for Kar2, the Hsp70 chaperone acting at the ER lumen (Gardner *et al*, 2013). On the other hand, both heat and salt stress have been shown to inhibit the G1/S regulon (Rowley *et al*, 1993; Bellí *et al*, 2001; Trotter *et al*, 2001). Thus, we reasoned that chaperone titration by stress could effectively reduce chaperone availability and restrain nuclear accumulation of Cln3 in a temporary manner (Fig 5A). We first interrogated our model and simulated a stress event by transferring different fractions of the folded protein (Prot_F_) to the unfolded population (ProtU). As shown in Fig 5B, the model predicted a sharp and transient reduction in available Ydj1 (Ydj1_A_) and free Cln3 (Cln3_F_). Notably, we found that Ydj1-GFP diffusion decreased very rapidly under both heat and salt stress (Fig 5C and D), and recovered at later times to attain similar steady-states to the pre-stress situation. Moreover, nuclear levels of Cln3-3HA displayed a similar and transient decrease after heat and salt stress (Fig 5E and F), which did not affect overall Cln3-3HA levels as measured by immunoblotting. Due to its extremely short half-life, Cln3 is thought to respond very rapidly to new conditions. However, nuclear levels of Cln3 only recovered pre-stress values after 20-30 min (Fig 5G and H), within the same time range needed by the Ydj1 chaperone to recover pre-stress mobility (Fig 5C and D), thus reinforcing a functional link between chaperone availability and nuclear accumulation of Cln3 under stress conditions. To test this further, we used our model to predict the behavior of Cln3 during stress adaptation assuming that overall protein folding and nuclear accumulation of Cln3 use chaperones in independent or competing manners. As seen in Fig 5G and H, only the competition scenario was able to recapitulate the Cln3 immunofluorescence data from both heat and salt stresses.

**Figure 5.**
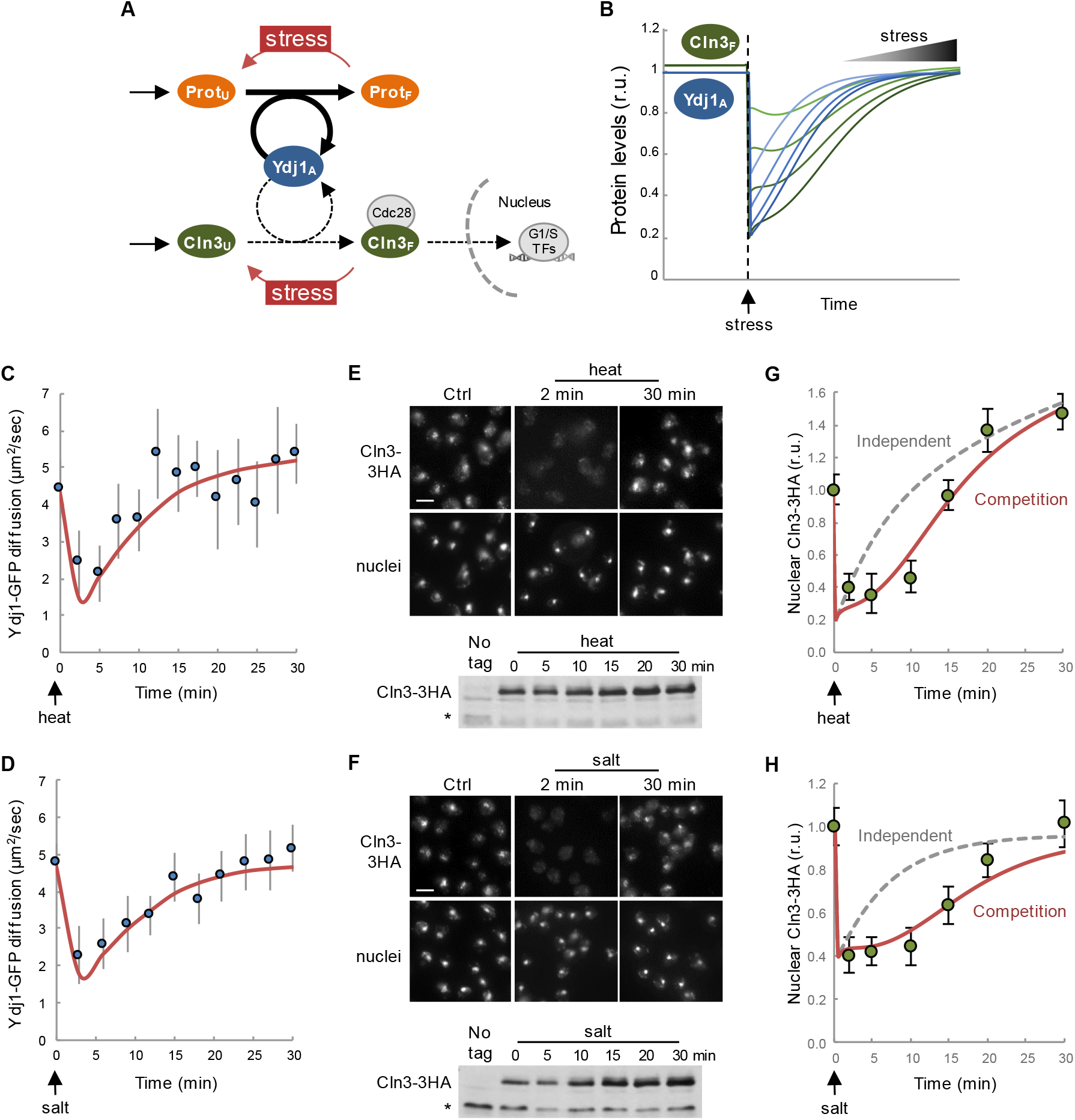
Stress reduces chaperone mobility and hinders nuclear accumulation of Cln3. A The competition model predicts that a sudden increase in the level of unfolded proteins (ProtU) would decrease the level of available chaperone (Ydj1_A_) and, hence, reduce the level of free Cln3 (Cln3_F_). B Prediction of available chaperone (Ydj1_A_) and free Cln3 (Cln3_F_) levels as a function of time after stress. Simulations were produced by using parameter set 3114 and transferring different fractions (20-80%) of folded protein (Prot_F_) to the unfolded population (ProtU). C,D Mean Ydj1-GFP diffusion coefficients (N>20, filled circles) and confidence limits (α=0.05, thin lines) are plotted at different times during heat (C) or salt (D) stress. Model fits (see text for details) are also shown (orange lines). E, F Immunofluorescence of Cln3-3HA in late-G1 cells arrested with α factor during heat (E) and salt (F) stress. Bottom: Cln3-3HA levels by immunoblotting. A cross-reacting band is shown as loading control. G,H Nuclear levels of Cln3-3HA quantified from cells during heat (E) and salt (F) stress as in panels E and F. Relative mean values (N>200) and confidence limits (α=0.05, thin horizontal lines) for the mean are shown. Model simulations (see text for details) assuming that ProtU and Cln3U require Ydj1_A_ in independent (gray dashed lines) or competing (solid orange lines) scenarios are also shown.

ER stress causes protein aggregation in the cytoplasm (Hamdan *et al*, 2017), and increases Ssa4 expression to levels very similar to Kar2 (Travers *et al*, 2000), suggesting that ER stress also affects Ydj1 availability. Moreover, ER stress also causes a G1 delay (Vai *et al*, 1987). We found that, similar to heat and salt stress, addition of tunicamycin to induce irreversible ER stress in yeast cells decreased Ydj1 mobility and nuclear levels of Cln3-3HA (Fig EV8). Overall, these data indicate that stress conditions due to very different environmental cues cause coincident decreases in Ydj1 mobility and Cln3 nuclear accumulation, reinforcing the notion that chaperone availability is a key parameter that controls G1 cyclin fate and adapts cell-cycle entry to growth and stress.

## DISCUSSION

Our data show that multifunctional chaperone systems play a limiting role in driving nuclear accumulation of the Cdc28-Cln3 complex during G1 progression. We also show that increased growth rates, by allocating higher levels of chaperones to protein synthesis and trafficking, reduce free chaperone pools and restrain nuclear accumulation of Cln3 at a given volume in G1. If Cln3 has to reach a threshold concentration to execute Start (Wang *et al*, 2009; Schmoller *et al*, 2015; Liu *et al*, 2015), fast growing cells would delay G1 progression to grow larger and attain the same pool of free chaperones and nuclear Cln3 (Fig 6). As a consequence, the critical size would be set as a function of growth rate. Due to the close molecular connection proposed here between protein synthesis and folding/release of the G1 cyclin, our model would allow cells to quickly adjust their size to many different intrinsic and extrinsic signals as long as they modulate protein synthesis, thus acting as a common mediator of specific growth signaling pathways and the Start network. This view would also explain, at least in part, why deleterious mutations in nutrient sensing pathways that control ribosome biogenesis and cell growth cause a small cell size phenotype (Jorgensen *et al*, 2004; Baroni *et al*, 1994; Tokiwa *et al*, 1994). A recent comprehensive analysis of cell size mutants concluded that, in the majority of *whi* mutants, the small cell size was due to indirect effects mostly caused by a decrease in growth rate (Soifer & Barkai, 2014).

**Figure 6.**
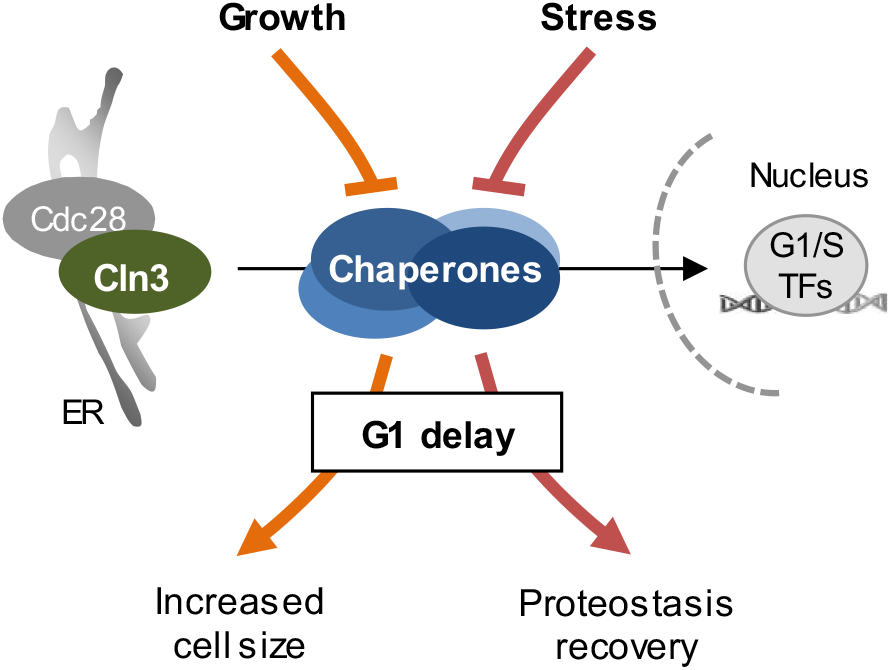
Competition for chaperones and the regulation of cell-cycle entry by growth and stress. Chaperone-dependent Cln3 folding and release would act as a key modulator of Start to subordinate cell-cycle entry to growth and stress. By compromising higher levels of chaperones in growth processes, fast growing cells would restrain nuclear accumulation of Cln3 until a larger cell size is attained. On the other hand, stressful conditions would compromise common chaperones and delay Start until normal proteostasis conditions are restored.

The chaperone competition mechanism would only operate above a minimal translation rate to sustain G1-cyclin synthesis (Schneider *et al*, 2004), and would collaborate with pathways regulating G1-cyclin expression by specific nutrients (Baroni *et al*, 1994; Newcomb *et al*, 2002; Gallego *et al*, 1997; Tokiwa *et al*, 1994). In this respect, Cln3 synthesis is strongly downregulated at low growth rates by a uORF-dependent mechanism (Polymenis & Schmidt, 1997), shifting cells from poor to rich carbon sources increases Cln3 abundance but yet delays Start in terms of cell size (Johnston *et al*, 1979; Hall *et al*, 1998; Tokiwa *et al*, 1994), phosphate deprivation increases degradation of Cln3 by Pho85-dependent phosphorylation (Menoyo *et al*, 2013), and Rim15 counteracts Whi5 dephosphorylation in G1 to facilitate cell-cycle entry in poor carbon sources (Talarek *et al*, 2017). Moreover, nutrient modulation of cell size is largely independent of the core components of the Start network (Jorgensen *et al*, 2004). From this point of view, modulation of the critical cell size threshold by nutrients and growth rate has remained a mysterious issue during decades (Johnston *et al*, 1979; Tyson *et al*, 1979; Fantes & Nurse, 1977). Our findings provide a general mechanism to modulate nuclear accumulation of Cln3 and cell-cycle entry as a function of growth rate.

Cells deficient for Cln3 still maintain a growth-rate dependent size at Start, but with a much larger variability compared to wild type cells (Yahya *et al*, 2014). Cln1 and Cln2 become essential in the absence of Cln3, and cells lacking these two other G1 cyclins also display a wider range of sizes at Start as a function of growth rate. Moreover, the *CLN2* mRNA has also been found enriched in Whi3 pulldowns (Colomina *et al*, 2008; Holmes *et al*, 2013), suggesting that the chaperone competition mechanism would also apply to Cln2, particularly in the absence of Cln3.

Whi5 levels have been found to decrease in rich carbon sources (Liu *et al*, 2015). If these modulation was responsible for cell size adaptation, we should expect cells displaying the opposite behavior, i.e. cells should be smaller in rich carbon sources. However, this observation could be interpreted as an effect, rather than a causative determinant of cell size adaptation. Whi5 is synthesized in a size-independent manner (Schmoller *et al*, 2015) and, also likely, in a growth-rate independent manner. As cells growing in rich carbon sources are larger, this effect would produce a decrease in Whi5 concentration. Thus, while Whi5 would act as a growth rate independent sizer, chaperone availability would dynamically transmit growth rate information to modulate G1 Cdk activation and adjust cell size as a function of the individual growth potential (Fig 6).

Available models of the cell cycle of budding yeast have centered their attention to different aspects of the molecular machineries that execute and regulate key transitions, but the implications of growth rate on cell size have been addressed only in a few occasions. Thus, a model based on intrinsic size homeostasis predicts growth-rate dependence if nutrient uptake is subject to geometric constraints (Spiesser *et al*, 2012, 2015). In this regard, surface to volume ratios have been recently highlighted in bacteria as key parameters for setting cell size as a function of growth rate (Harris & Theriot, 2016). Interestingly, an important fraction of Ydj1 is involved in post-translational protein translocation at the ER (Caplan *et al*, 1992; McClellan *et al*, 1998) to fuel cell surface growth which, as a consequence, would set Ydj1 demands as a function of surface-to-volume ratios. On the other hand, in a recent model of the G1/S transition, it has been proposed that growth-rate dependence would be exerted, directly or indirectly, by ribosome-biogenesis effects on the Cln3/Whi5 interplay (Palumbo *et al*, 2016). In agreement with this idea, here we propose that chaperones act as key mediators in this pathway by modulating the ability of Cln3 to accumulate in the nucleus and, hence, attain a critical Cln3/Whi5 ratio.

Nutrient modulation of cell size also takes place during bud growth and, as we had observed in G1, cell size at birth displays a strong correlation with growth rate at a single cell level during the preceding S-G2-M phases (Leitao & Kellogg, 2017). Thus, mechanisms sensing growth rate independently of the specific nutritional conditions also operate during bud volume growth.

Finally, we show that different stress agents cause concurrent decreases in Ydj1 mobility and nuclear Cln3 levels, supporting the idea that chaperone availability is a key factor controlling localization of the most upstream G1 cyclin and, hence, modulating G1 length under stress conditions. Heat shock transiently inhibits G1/S gene expression (Rowley *et al*, 1993), but the molecular mechanism is still unknown. On the other hand, osmotic shock causes a similar temporary repression of the G1/S regulon (Bellí *et al*, 2001), where Hog1-mediated phosphorylation of Whi5 and Msa1 plays an important role in transcription inhibition (González-Novo *et al*, 2015). However, G1/S gene expression was still repressed by salt in the absence of Whi5 and Msa1, indicating the existence of additional mechanisms sufficient to inhibit the G1/S regulon under osmotic shock. it has long been known that ER stress causes a G1 delay (Vai *et al*, 1987), but the molecular mechanisms have not been described. Our results uncover a new mechanism that would explain the observed downregulation of G1/S genes by heat, salt and ER stress, whereby immediate titration of chaperones would decrease available pools of Ssa1 and Ydj1 chaperones required to accumulate the G1 Cdk in the nucleus for triggering Start. In this manner, stress conditions would delay G1 progression until proteostasis and chaperone availability recover to normal levels (Fig 6).

Free chaperone levels could also report growth capability to other processes influenced by growth rate (Brauer *et al*, 2008) or stressful conditions (De Nadal *et al*, 2011) through similar competition settings. Furthermore, imposing a high level of free chaperones as a requirement for Start would ensure their availability in highly-demanding downstream processes such as polarized growth for bud emergence or nucleosome remodeling during replication.

## MATERIALS AND METHODS

### Strains and plasmids

Yeast strains and plasmids used are listed in Supplementary Table 1. Parental strains and methods used for chromosomal gene transplacement and PCR-based directed mutagenesis have been described (Gallego *et al*, 1997; Ferrezuelo *et al*, 2012). Unless stated otherwise, all gene fusions in this study were expressed at endogenous levels at their respective loci. As C-terminal fusions of GFP or other tags has strong deleterious effects on Ydj1 function (Saarikangas *et al*, 2017), we inserted GFP at amino acid 387, between the dimerization domain and the C-terminal farnesylation sequence of Ydj1. This construct had no detectable effects on growth rate or cell volume when expressed at endogenous levels. The mCitrine-Cln3-11A fusion protein contains a hypoactive and hyperstable cyclin with 11 amino acid substitutions (R108A, T420A, S449A, T455A, S462A, S464A, S468A, T478A, S514A, T517A, T520A) that allows its detection by fluorescence microscopy with no gross effects on cell cycle progression (Schmoller *et al*, 2015).

### Growth conditions

Cells were grown for 7-8 generations in SC medium with 2% glucose at 30°C before microscopy unless stated otherwise. Other carbon sources used were 2% galactose, 2% raffinose and 3% ethanol. GAL1p-driven gene expression was induced by addition of 2% galactose to cultures grown in 2% raffinose at OD600=0.5. When stated, 10 μM β-estradiol was used to induce the GAL1 promoter in strains expressing the Gal4-hER-VP16 (GEV) transactivator (Louvion *et al*, 1993). Azetidine 2-carboxylic acid (AZC) was used at 10 mM, and cycloheximide was added at a sublethal dose of 0.2 μg/ml that does not trigger stress gene activation (Jacquet, 2003; Trotter *et al*, 2002). Heat and osmotic stresses were imposed by transferring cells from 25ºC to 37ºC or adding 0.4M NaCl at 30ºC, respectively. Tunicamycin was added used at 1 μg/ml. Small newly-born cells were isolated from Ficoll gradients (Mitchison, 1988).

### Time-lapse microscopy

Cells were analyzed by time-lapse microscopy in 35-mm glass-bottom culture dishes (GWST-3522, WillCo) in SC-based media at 30°C essentially as described (Ferrezuelo *et al*, 2012) using a fully-motorized Leica AF7000 microscope. Time-lapse images were analyzed with the aid of BudJ (Ferrezuelo *et al*, 2012), an ImageJ (Wayne Rasband, NIH) plugin that can be obtained from www.ibmb.csic.es\home\maldea to obtain cell dimensions and fluorescence data. Volume growth rate in G1 were obtained as described (Ferrezuelo *et al*, 2012). Start was estimated at a single-cell level as the time where the nuclear to cytoplasmic ratio of Whi5 had decreased below 1.5. Photobleaching during acquisition was negligible (less than 0.1% per time point) and autofluorescence was always subtracted.

### Determination of nuclear and cellular concentrations of fluorescent fusion proteins

Wide-field microscopy is able to collect the total fluorescence emitted by yeast cells and, consequently, cellular concentration of fluorescent fusion proteins was obtained by dividing the integrated fluorescence signal within the projected area of the cell by its volume. Regarding the quantification of nuclear levels, since the signal in the nuclear projected area is influenced by both nuclear and cytoplasmic fluorescence, determination of the nuclear concentration required specific calculations as described in Fig EV1A. In confocal microscopy the fluorescence signal is directly proportional to the concentration of the fluorescent fusion protein, and required no further calculations. The nuclear compartment was delimited as described (Ferrezuelo *et al*, 2012).

### Chaperone mobility analysis by FLIP and FCS

We used fluorescence loss in photobleaching (FLIP) and fluorescence correlation spectroscopy (FCS) to analyze chaperone mobility in a Zeiss LSM780 confocal microscope. FLIP was used as a qualitative assay to determine Ssa1-GFP and Ydj1-GFP mobility in the whole cell. A small circular region of the cytoplasm (3.6 μm^2^) was repetitively photobleached at full laser power while the cell was imaged at low intensity every 0.5 sec to record fluorescence loss. After background subtraction, fluorescence data from an unbleached cell region were made relative to the initial time point, and a mobility index was calculated as the inverse of the fluorescence half-life obtained by fitting an exponential function to fluorescence emitted as a function of time. Quantitative analysis of Ydj1-GFP diffusion by FCS was performed essentially as described (Saarikangas *et al*, 2017). Specifically, FCS analysis was done at 25°C to minimize signal variability in the 0.1-1 sec range, and cells were prebleached to attain count rates within the 50-100 kHz range during acquisition for periods of 5 sec. FCS correlation data were fitted in the 10 μsec to 100 msec range of time intervals with the aid of QuickFit 3 (http://www.dkfz.de/Macromol/quickfit/), assuming a 1-component anomalous mode of diffusion (α=0.5) in the Levenberg-Marquardt algorithm to obtain diffusion coefficients. Duplicate measurements were always taken and outliers were removed from analysis if the relative standard error of the fitted coefficient of diffusion was higher than 50%, or the fitted autocorrelation intersect was higher than 1.01 as a result of strong perturbations in the average count rate during acquisition. In time-lapse experiments outliers were removed if the relative difference to neighbor values was higher than 50%. Removed outliers were always less than 5% of measurements.

### Nuclear import rate determinations by FLIP

To analyze nuclear import kinetics of Cdc28-GFP, a circle inscribed within the Htb2-mCherry nuclear region was repetitively photobleached while the cell was imaged every 0.5 sec to record fluorescence loss. After background subtraction, fluorescence data were corrected with those from a non-bleached cell. Finally, fluorescence signals within nuclear and cytoplasmic areas were used to analyze import kinetics as described in Fig EV2A. The export rate was assumed constant among G1 cells and obtained as described in Fig EV2B.

### Immunofluorescence

Immunofluorescence of endogenous levels of Cln3-3HA was done by a signal-amplification method (Vergés *et al*, 2007) with αHA (clone 3F10, Roche) and goat-αrat and rabbit-αgoat Alexa555-labeled antibodies (Molecular Probes) on methanol-pemeabilized cells. To analyze localization of Cln3-3HA we used N2CJ, a plugin for ImageJ (Wayne Rasband, NIH), to perform accurate quantification in both cytoplasmic and nuclear compartments of cells (Yahya *et al*, 2014).

### Model equations

The model in Fig 3A was simulated with a set of non-linear differential equations.

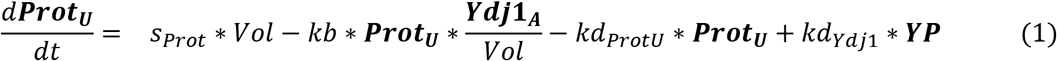

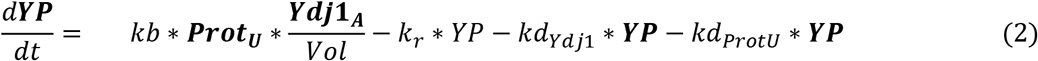

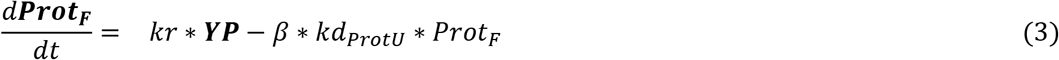

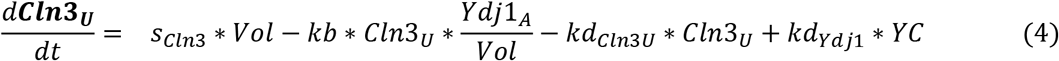

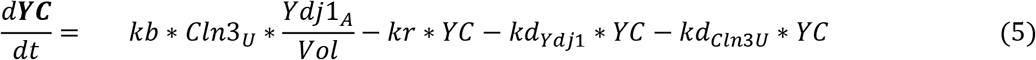

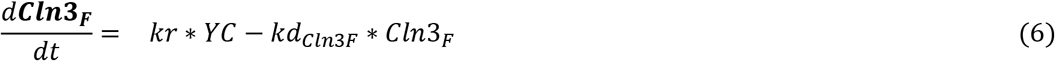

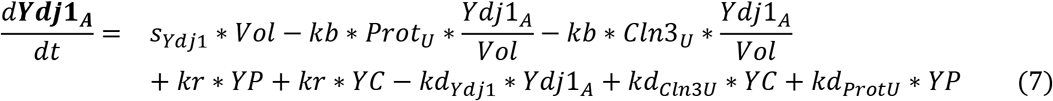

This model has seven state variables: ***Prot_U_***, unfolded proteins; ***YP***, Ydj1-bound proteins; ***Prot_F_***, folded proteins; ***Cln3_U_***, unfolded Cln3; YC, Ydj1-bound Cln3; ***Cln3_F_***, folded Cln3; and ***Ydj1_A_***, free available Ydj1.

### Model parameters and simulations

In the model we define eleven parameters (Table 1). Ydj1 binding and release (*kb, kr*), which are assumed to be the same for *Prot_U_* and *Cln3_U_* (see Fig EV4), and synthesis (*S_prot_, S_cin3_, S_ydji_*) and degradation (*kd_protU_, kd_cln3u_, kd_Cln3_F__, kd_Ydj1_*) of the different components. A scaling factor (β=0.01) is used to ensure that folded protein (*Prot_F_*) has a half-life 100 times that of unfolded protein (*Prot_U_*). The state variables in the model are in units of molecular number, not concentration, and therefore all 0^th^ and 2^nd^ order reactions are explicitly scaled by cell volume (*Vol*). In the model, we assume that the rate of change of cell volume with time is much lower than the rates of the biochemical reactions studied. This allowed us to treat the cell volume as a pseudo-parameter, so the steady state of other variables with respect to cell volume and growth rate (*S_protU_*) could be analyzed more straightforwardly. The half-life of unfolded (*kd_Cln3U_*) Cln3 was set to be 1.8 times longer than folded (*kd_Cln3_F__*) Cln3 as deduced from steady-state levels of Cln3-3HA in wild-type and Ydj1-deficient cells (Yaglom *et al*, 1996), while Ydj1 is an stable protein, with a half-life 20 times longer than folded Cln3. To reduce the degrees of freedom available for modeling, degradation rates of all molecules were kept constant independent if they are in a complex or free. The underlying reasoning is that molecules should be still recognized by their respective degradation machinery independent of their binding partners. This should be true in our cases, since the binding of Ydj1 to its targets is highly dynamic. We converted half-lives to degradation rates using the formula *λ* = log (2)/*t*_1/2_. The remaining five parameters were used to quantitatively fit two sets of measurements: the relationship between the diffusion rate of Ydj1 and growth rate in G1 cells (Fig 3F), and the relationship between budding volume and growth rate (Fig 3G).

**Table 1.** Model parameters

| Parameter | Definition | Type | Value | Parameter set 3114 |
| --- | --- | --- | --- | --- |
| $Vol$ | Cell volume | Controlled | 10-100 fl | |
| $S_{ProtU}$ | Protein synthesis | Controlled | 0.01-1 molec·sec<sup>-1</sup>·fl<sup>-1</sup> | |
| $S_{Cln3}$ | Cln3 synthesis | Fitted | 4.9·10<sup>-2</sup>-4.8 molec·sec<sup>-1</sup>·fl<sup>-1</sup> | 0.126 molec·sec<sup>-1</sup>·fl<sup>-1</sup> |
| $S_{Ydj1}$ | Ydj1 synthesis | Fitted | 4.8·10<sup>-5</sup>-6.0·10<sup>-2</sup> molec·sec<sup>-1</sup>·fl<sup>-1</sup> | 9.19·10<sup>-4</sup> molec·sec<sup>-1</sup>·fl<sup>-1</sup> |
| $kd_{ProtU}$ | ProtU degradation | Fitted | 6.4·10<sup>-3</sup>-31.5 sec<sup>-1</sup> | 7.26·10<sup>-2</sup> sec<sup>-1</sup> |
| $kd_{ProtF}$ | ProtF degradation | Fitted | 0.01* $kd_{ProtU}$ sec<sup>-1</sup> | 7.26·10<sup>-4</sup> sec<sup>-1</sup> |
| $kb$ | Binding to Ydj1 | Fitted | 1.4·10<sup>-2</sup>-26.7 fl·molec<sup>-1</sup>·sec<sup>-1</sup> | 0.891 fl·molec<sup>-1</sup>·sec<sup>-1</sup> |
| $kr$ | Release from Ydj1 | Fitted | 2.8·10<sup>-3</sup>-7.7 sec<sup>-1</sup> | 0.388 sec<sup>-1</sup> |
| $kd_{Cln3U}$ | Cln3U degradation | Fixed | 6.93·10<sup>-2</sup> sec<sup>-1</sup> | |
| $kd_{Cln3F}$ | Cln3F degradation | Fixed | 3.85·10<sup>-2</sup> sec<sup>-1</sup> | |
| $kd_{Ydj1}$ | Ydj1 degradation | Fixed | 5.77·10<sup>-3</sup> sec<sup>-1</sup> | |

### Budding volume *vs* growth rate

In the model, we assume that Start is triggered by Cln3 when reaches a given critical threshold 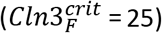 that, together with Cdc28, is needed to phosphorylate and inactivate a fixed given amount of DNA-bound Whi5/SBF complexes (Schmoller *et al*, 2015; Wang *et al*, 2009). Our objective was then to minimize the difference between experimental measurements and simulation results by changing the five free parameters in *L*1 (*θ*) = (*Vol* (*S_Vol_*)*_exp_* – *Vol* (*θ* | *S_Vol_*)*_sim_*)^2^, where *Vol* (*S_Vol_*)*_exp_* is the experimentally measured budding volume (as a proxy for the critical volume at Start) of a cell growing at rate *S_Vol_*, and *Vol* (*θ* | *S_Vol_*)*_sim_* is the volume of cells growing at rate *S_Vol_* reach a steady state value where *Cln3_F_* equals the critical threshold 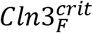.

### Ydj1 diffusion *vs* growth rate

We assume that Ydj1 is present in two distinct pools: a fast diffusing free fraction (*Ydj1_A_*), and a slowly diffusing fraction bound to either *Prot* or *Cln3* (*YP+YC*). To compare the experimentally measured diffusion rate of Ydj1 to our simulations, we define the simulated diffusion rate of Ydj1 as the weighted average diffusion coefficient for all species of Ydj1 in our model as follows:

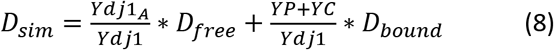

For Ydj1-GFP we estimated that *D_bound_* = 1 μm^2^/min from minimal values observed in AZC-treated cells, and *D_free_* = 30 μm^2^/min from maximal values obtained with GFP and corrected by the different Stokes radius. As above, we sought to minimize 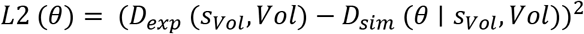, where *D_exp_* (*s_Vol_*, *Vol*) and *D_sim_*(*θ* | *s_Vol_*, *Vol*) are the experimental and simulated diffusion coefficients at a given growth rate and volume in G1 cells.

### Fitting procedures

For fitting Ydj1 diffusion and budding volume as a function of growth rate we combined the respective square differences to seek 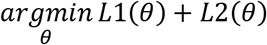. To find the optimal value of *θ* we used a variant of the Levenberg-Marquardt algorithm with a 2^nd^ order correction (Transtrum & Sethna, 2012). As all the simulations were done at steady state, to obtain the Jacobian of the steady state with respect to the parameters, we applied the implicit function theorem following the approach described here: https://arxiv.org/pdf/1602.02355.pdf. Numerical integration of the differential equations (to obtain an initial value for the steady state root finding) was done using the *SciPy routine odeint*, which automatically switches between stiff and non-stiff solvers. After identifying the minima, we performed a Markov Chain Monte Carlo (Goodman & Weare, 2010) exploration of the parameter space using *emcee* (Foreman-Mackey *et al*, 2013). After discarding the initial 3000 draws of the MC chain as burn-in, which were too close to the minima, we collected 1500 sets with good match to data and repeated all simulations using this ensemble of parameters to estimate the uncertainty in parameters (Fig 3B) and simulations of variables used in the initial exhaustive fitting process (Fig 3F and 3G). Simulations of other variables that were experimentally tested (Fig 3H, 4B and 5B) were done with parameter set 3114 (Table 1), which gave the minimal p value in the fitting process, and using a 4-fold range of values for the Ydj1-binding and release constant rates (*kb* and *kr*, respectively). CHX effects were simulated by reducing *ksProt, ksCln3* and *ksYdj1* 3, 4.5, 6, 9, or 12-fold to test different conditions around the experimental reduction of 5.9-fold (Fig EV7). Stress effects were simulated on steady states by transferring different fractions (20-80%) of the folded protein (Prot_F_ and Cln3_F_) to the unfolded population (ProtU and Cln3U, respectively). To simulate transient effects in Cln3_F_ by stress, experimental data of Ydj1-GFP mobility were used to fit the model time variable at different percentages of protein unfolding (Fig 5CD), resulting in 60% for salt stress and 80% for heat stress. We took into account that heat, but not salt, stress increases Ydj1 mRNA levels by 2.9-fold (Gasch *et al*, 2000). Then, the resulting fitted parameters were applied to the model to simulate the evolution of Cln3_F_ after stress assuming that ProtU and Cln3U compete or not for Ydj1_A_ (Fig 5GH). The model was deposited in the BioModels (Chelliah *et al*, 2015) database as MODEL1808310001 in SBML format and a Copasi (Hoops *et al*, 2006) file to reproduce simulations with parameter set 3114. Code of the estimation of minima in IPython Notebooks is available upon request.

### Parameter distributions

As amply described in the literature, parameters in systems biology models are disheveled (Gutenkunst *et al*, 2007) or structurally unidentifiable (Szederkényi *et al*, 2011). We observed this behavior as well (Fig 3B), with parameters consistent with the experimental data spanning several orders of magnitude.

### Statistical analysis

We routinely show the standard error of the mean (N<10) or confidence limits at α=0.05 (N>10) to allow direct evaluation of variability and differences between mean values in plots. Sample size is always indicated in the figure legend and, when appropriate, t-test p values are shown in the text. For model predictions the mean values and standard deviations are plotted. All experiments were done at least twice with fully independent cell samples.

### Miscellaneous

*In vitro* luciferase refolding assays have been described (Summers *et al*, 2009). Translational efficiency of refolding extracts was measured by incubation with 0.1 mM puromycin. Immunoblot analysis with αpuromycin (clone 12D10, Sigma), αHA (clone 12CA5, Roche) and αDpm1 (clone 5C5, Molecular Probes) was as described (Georgieva *et al*, 2015). Ydj1 binding assays to GST-fusions of Cln3, luciferase and P6, a selected Ydj1-target peptide (Kota *et al*, 2009) used as reference, were done with purified proteins as described (Lee *et al*, 2002). Protein synthesis rates in live cells were determined by S^35^-methionine incorporation (Gallego *et al*, 1997).

## ACKNOWLEDGEMENTS

We thank A. Cornadó and E. Rebollo for technical assistance, T.Zimmermann for technical advice in FCS experiments, and B. Futcher and J. Skotheim for providing strains. We also thank C. Rose for editing the manuscript, and F. Antequera, Y. Barral, C. Gallego, J.C. Igual, S. Oliferenko and F. Posas for helpful comments. This work was funded by the Spanish Ministry of Science, Consolider-Ingenio 2010, and the European Union (FEDER) to M.A. D.F.M. received a FI fellowship from *Generalitat de Catalunya*.

## AUTHOR CONTRIBUTIONS

D.F.M., E.P. and G.Y. built genetic constructs and strains, and performed the experiments. F.V. and A. C.- N. implemented the mathematical model and performed the informatic analysis. A. C.-N. and M.A. conceived the study, designed the mathematical model, analyzed the data and wrote the manuscript.

---

**Figure EV1.**
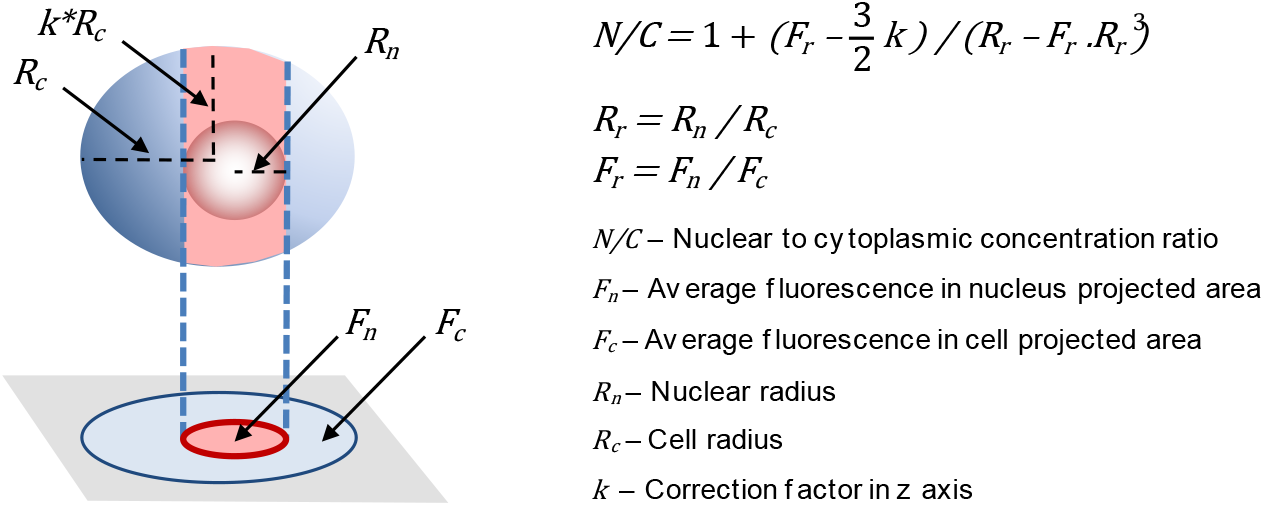
Nuclear to cytoplasmic ratios by wide-field fluorescence microscopy. Fluorescence within the nuclear projected area is influenced by fluorescence levels in both nuclear and cytoplasmic compartments. Briefly, the nuclear projection collects fluorescence originated from a column defined by the nuclear diameter and the cell height. Total fluorescence in the column is the weighted sum of nuclear and cytoplasmic concentrations of fluorophore assuming that the column is a cylinder and the nucleus a sphere with known dimensions relative to the cell. Cytoplasmic fluorescence (and concentration) is obtained from cell regions outside the nuclear projection, and the ratio of the nuclear to cytoplasmic concentration of the fluorescent protein is calculated as 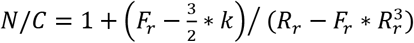, where *Fr* is the ratio of average fluorescence signal in nuclear (*F_n_*) and cell (*F_c_*) projected areas, *R_r_* is the ratio of nuclear (*R_n_*) and cell (*R_c_*) radius, and *k* corrects for cell radius variation in the z axis, which was experimentally obtained from GFP-expressing wild-type cells.

**Figure EV2.**
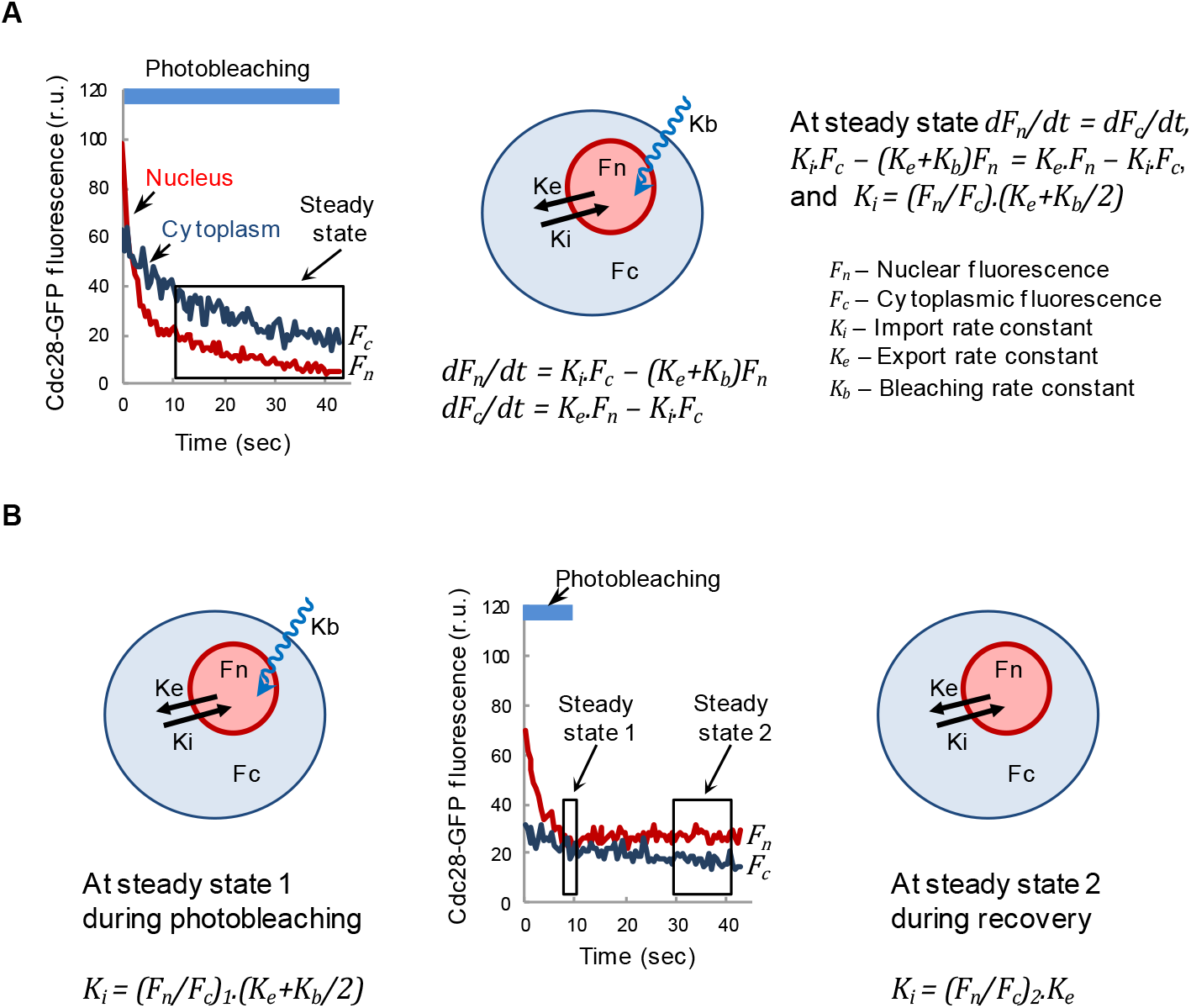
Nuclear import kinetics analysis by nuclear FLIP. A Schematic showing equations and parameters used to analyze Cdc28-GFP import and export kinetics as described under Materials and Methods. Fluorescence signals within nuclear (*F_n_*) and cytoplasmic (*F_c_*) areas were used to obtain a nuclear to cytoplasmic ratio (*F_n_*/*F_c_*). This ratio dropped to different extents during the first 10 sec of exposure, reflecting a population of Cdc28-GFP molecules in the nucleus not in dynamic equilibrium with the cytoplasm. After this period, however, the (*F_n_*/*F_c_*) ratio reached a steady state defined by *dF_n_/dt=dF_c_/dt*,being *d*F_n_*/dt = Ki.F_c_ – (K_e_+K_b_)*F_n_**, and *dF_c_/dt = K_e_.*F_n_* – K_i_.F_c_*,where *K_i_, K_e_* and *K_b_* are the import, export and bleaching rate constants, respectively. Thus, *K_i_.F_c_ – (K_e_+K_b_)*F_n_* = K_e_.*F_n_* – K_i_.F_c_*, and *K_i_ = (*F_n_*/F_c_) (Ke+Kb/2*). The bleaching rate constant was obtained from Htb2-mCherry fluorescence loss in the same cell and corrected by a bleaching factor of 1.94±0.09 to account for intrinsic differences in bleaching rates for GFP and mCherry, which was obtained from G1 cells (N=33) under conditions that minimize import contribution (Cln3-deficient cells). B Schematic showing equations and parameters used to obtain Cdc28-GFP export kinetics. In this case, the nuclear region of wild-type G1 cells (N=34) was only photobleached for 10 s and allowed to recover fluorescence by nuclear import of Cdc28-GFP. *F_n_/F_c_* ratio steady states were used to calculate import rate constants at photobleaching, *K_i_ = (F_n_/F_c_)1. (K_e_+K_b_/2)*, and recovery, *Ki = (F_n_/F_c_)2 .K_e_*, to finally obtain the export rate constant *K_e_* = 0.199±0.020.

**Figure EV3.**
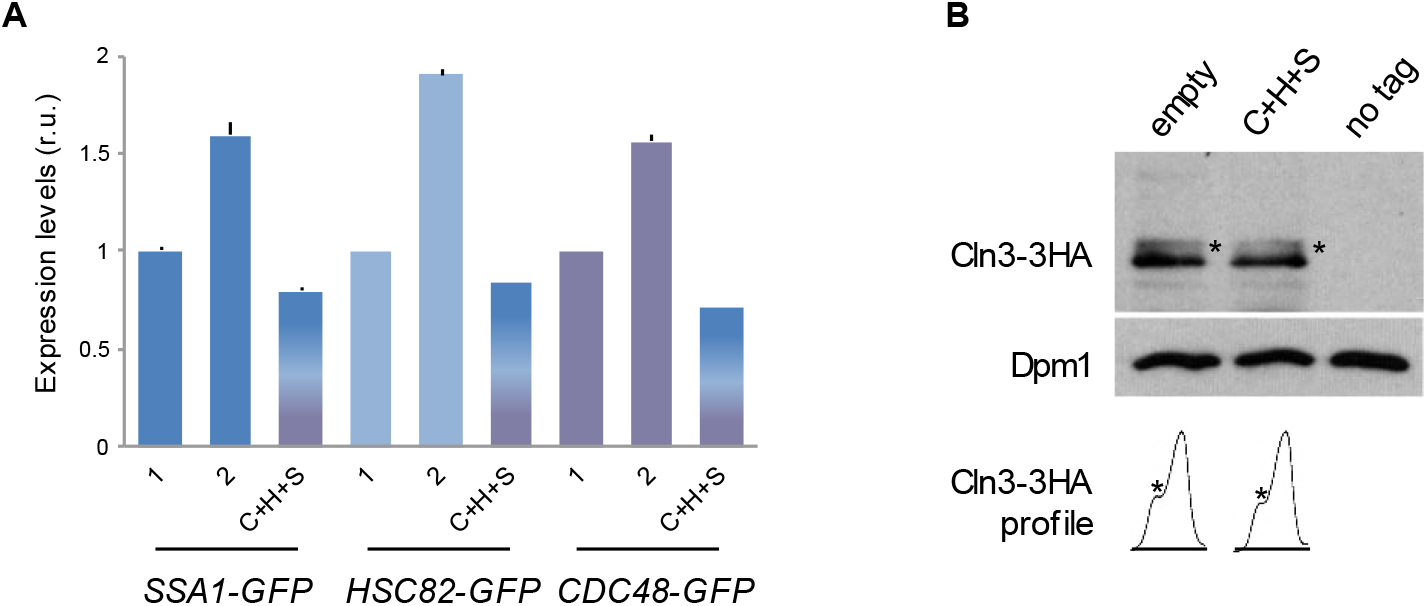
Gene dosage and expression levels of GFP-fused chaperones and Cln3-3HA. A Expression levels of the indicated chaperones endogenously expressed as GFP fusions in single (1) or double (2) copy, or in single copy and the presence of untagged-chaperone expressing vectors (C+H+S) as in Fig 2A. Fluorescence levels were determined and made relative to the mean value for cells containing a single copy of the indicated chaperone gene. Mean values (N>500) and confidence limits (α=0.05, thin vertical lines) for the mean are plotted. B Immunoblot analysis of endogenously expressed Cln3-3HA levels from cells containing empty or chaperone expressing vectors as in Fig 2A. A Dpm1 immunoblot is shown as loading control. Densitometric profiles of the corresponding lanes are shown for direct assessment of total levels and phosphorylated (*) status.

**Figure EV4.**
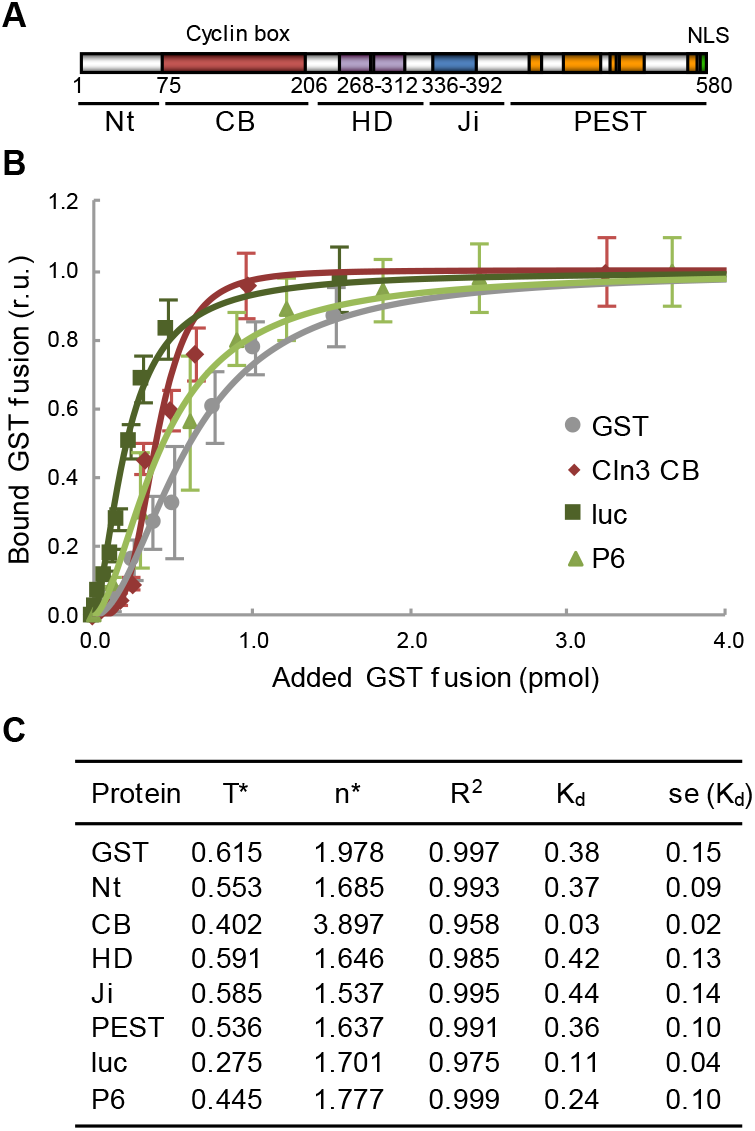
Ydj1 binding affinity to Cln3 and reference proteins. A Cln3 scheme indicating the N-terminal domain (N_t_), the cyclin box (CB), a hydrophobic bi-partite domain (HD), the inhibitory J domain (Ji), the C-terminal PEST rich region and the nuclear localization signal (NLS). B Binding efficiencies of increasing amounts (0-4 pmol) of the referred GST fusions to Ydj1 (1 pmol). Luciferase (luc) and P6, a selected Ydj1-target peptide, were used as references. Mean relative bound levels (N=3) and standard deviations are plotted. C Binding parameters of Ydj1 to the referred GST fusions in assays as in panel B. A simultaneous mode of binding was assumed for fitting the Hill equation, y = x^n^ / (T^n^+ x^n^). Obtained parameters and the resulting dissociation constant (K_d_ = T^n^) are shown.

**Figure EV5.**
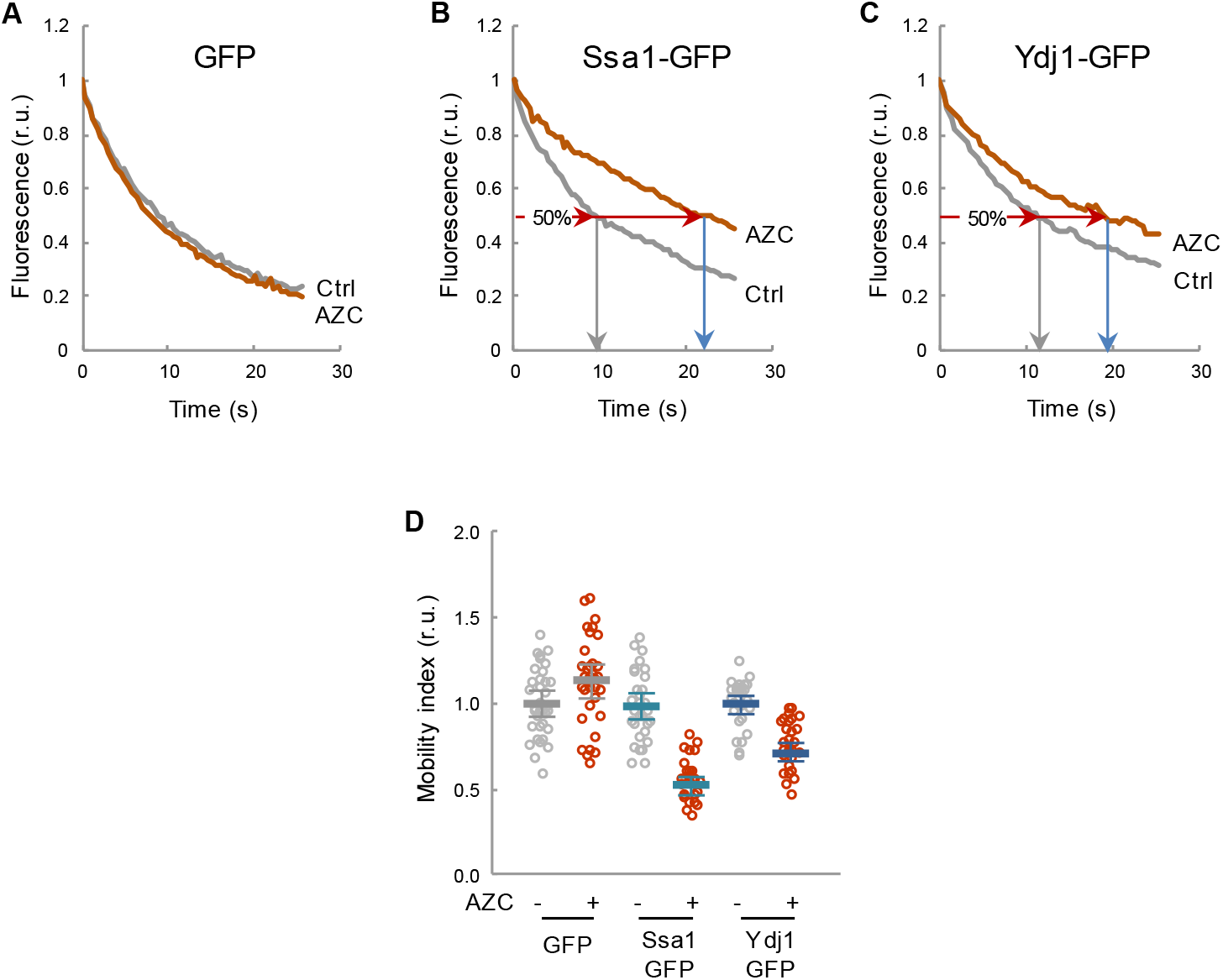
Mobility of GFP-fusion proteins as a reporter of chaperone availability. A-C Yeast cells expressing the indicated GFP fusions were analyzed by confocal microscopy to quantify intracellular mobility of Ssa1 and Ydj1 chaperones in the presence or absence of AZC. Since AZC-containing proteins accumulate with chaperones into disperse cellular aggregates (Escusa-Toret *et al*, 2013), FLIP was used to include both soluble and aggregated forms of these chaperones in the analysis. Representative fluorescence decay curves of individual cells are plotted. D Individual protein mobility values (N>25) of Ssa1- and Ydj1-GFP fusions obtained by FLIP before (-) or 15 min after adding AZC (+) are plotted. Mean values (thick lines) and confidence limits for the mean (α=0.05, thin lines) are also shown.

**Figure EV6.**
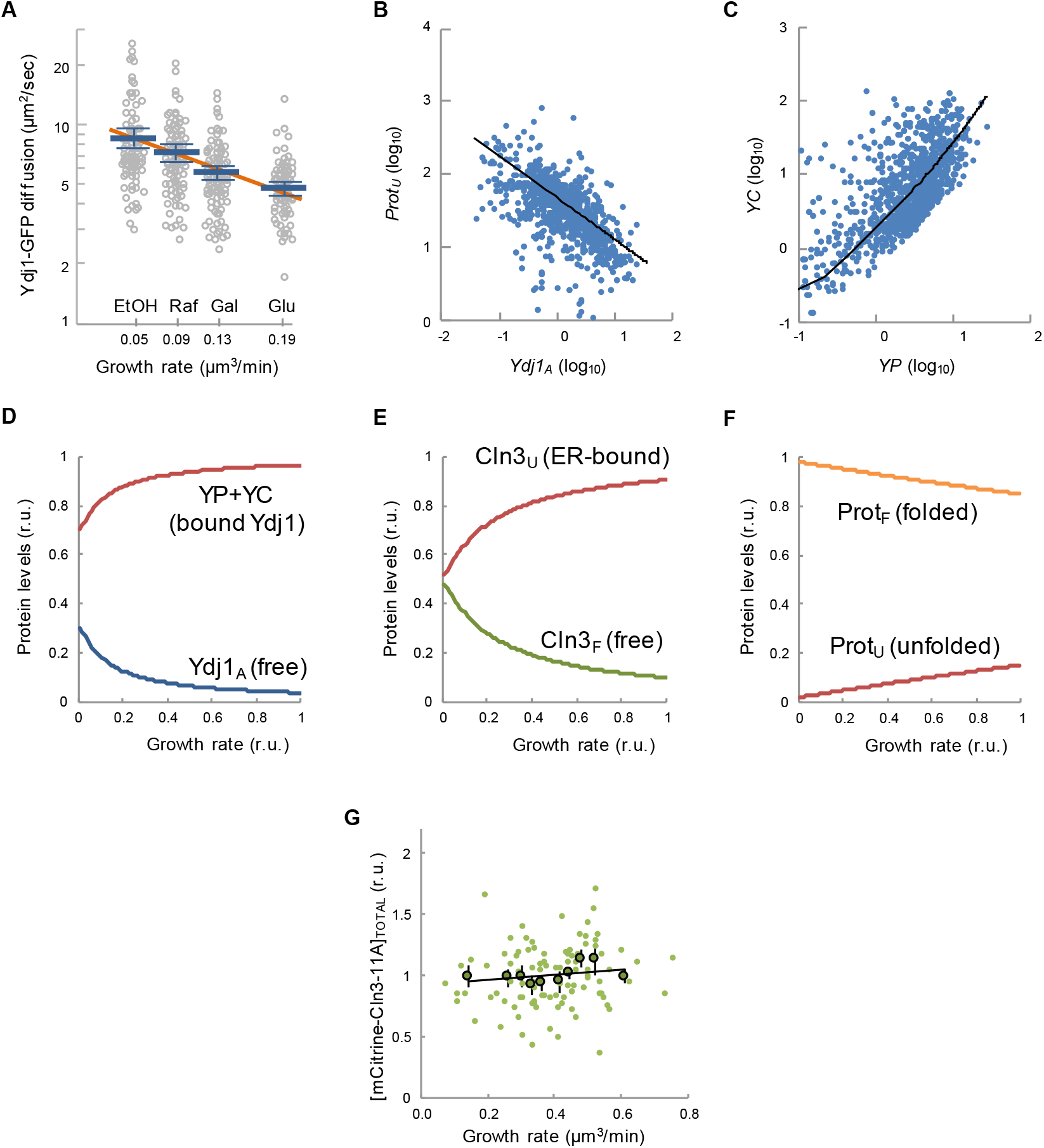
Ydj1-GFP mobility as a function of growth rate and correlation of state variables. A Ydj1-GFP diffusion assayed by FCS in cells growing at different population growth rates in the indicated carbon sources. Individual diffusion coefficients, mean values (N=90, thick blue lines), confidence limits for the mean (α=0.05, thin lines), and a linear fit (thick orange line) are plotted. B-C Covariation plots of *Prot_U_* vs *Ydj1_A_* (C), and *YC* vs *YP* (D) are shown. Values were obtained from parameters sets in Fig 3B and plotted in log space with the corresponding regression lines. D-F Prediction of main variables of the model as a function of growth rate. Parameter set 3114 was used to simulate relative levels of protein-bound (*YC* and *YP*) and available (*Ydj1_A_*) chaperone (D), unfolded ER-bound (*Cln3_U_*) and free folded (*Cln3_F_*) G1 cyclin (E), and all other unfolded (*Prot_U_*) and folded (*Prot_F_*) proteins (F). G Cells (N=100) were analyzed to obtain the mean cellular concentration of mCitrine-Cln3-11A during G1. Singl-cell (small circles) or binned data (large circles, 10 cells/bin) by growth rate are plotted. Confidence limits (α=0.05) and a linear regression line for binned data are also plotted.

**Figure EV7.**
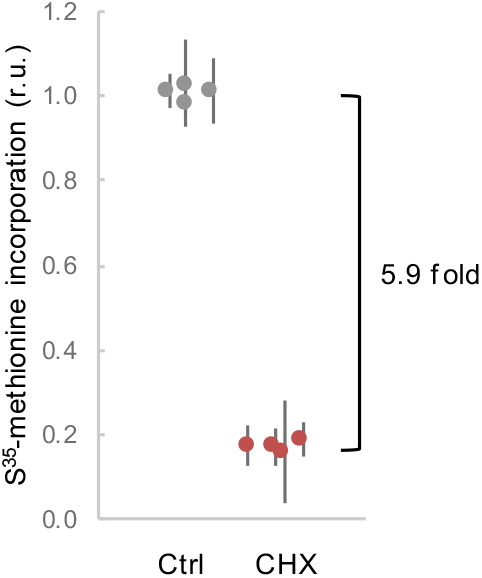
Protein synthesis effects by CHX at a sublethal dose. Protein synthesis as measured by S^35^-methionine incorporation in live cells in the absence or presence of 0.2 μg/ml CHX. Relative mean values (circles) of triplicate measurements from independent cell batches (N=4) and confidence limits for the mean (α=0.05, thin lines) are plotted.

**Figure EV8.**
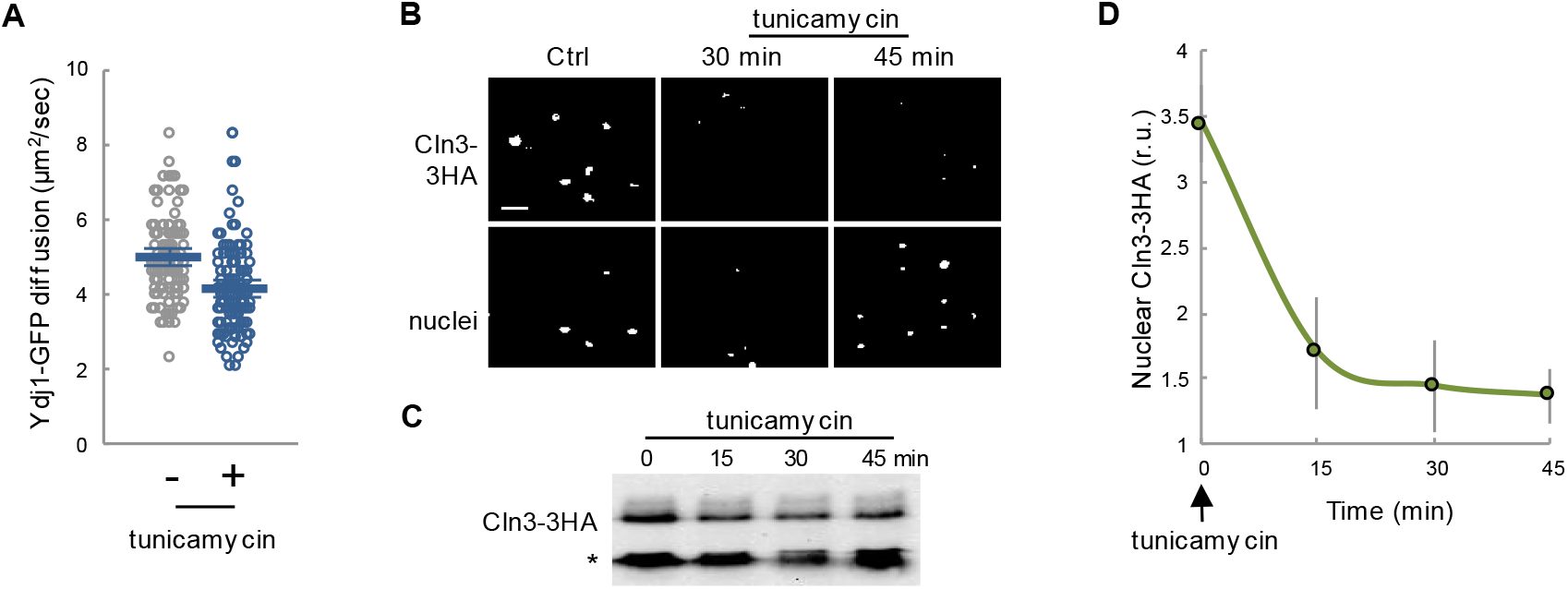
Effects of ER stress on Ydj1-GFP mobility and Cln3-3HA nuclear levels. A Yeast cells expressing Ydj1-GFP were analyzed by FCS 30 min after adding tunicamycin to 1 μg/ml. Individual protein diffusion coefficients are plotted (N>20). Mean values (thick lines) and confidence limits for the mean (α=0.05, thin lines) are also shown. B Immunofluorescence of Cln3-3HA in late-G1 cells arrested with α factor 30 min after 1 μg/ml tunicamycin addition. Bottom: Cln3-3HA levels by immunoblotting at different times in the presence of tunicamycin. A cross-reacting band is shown as loading control. C Nuclear levels of Cln3-3HA quantified from cells during treatment with tunicamycin as in panel B. Relative mean values (N>200) and confidence limits (α=0.05, thin horizontal lines) for the mean are shown.

**Supplementary Table 1.**
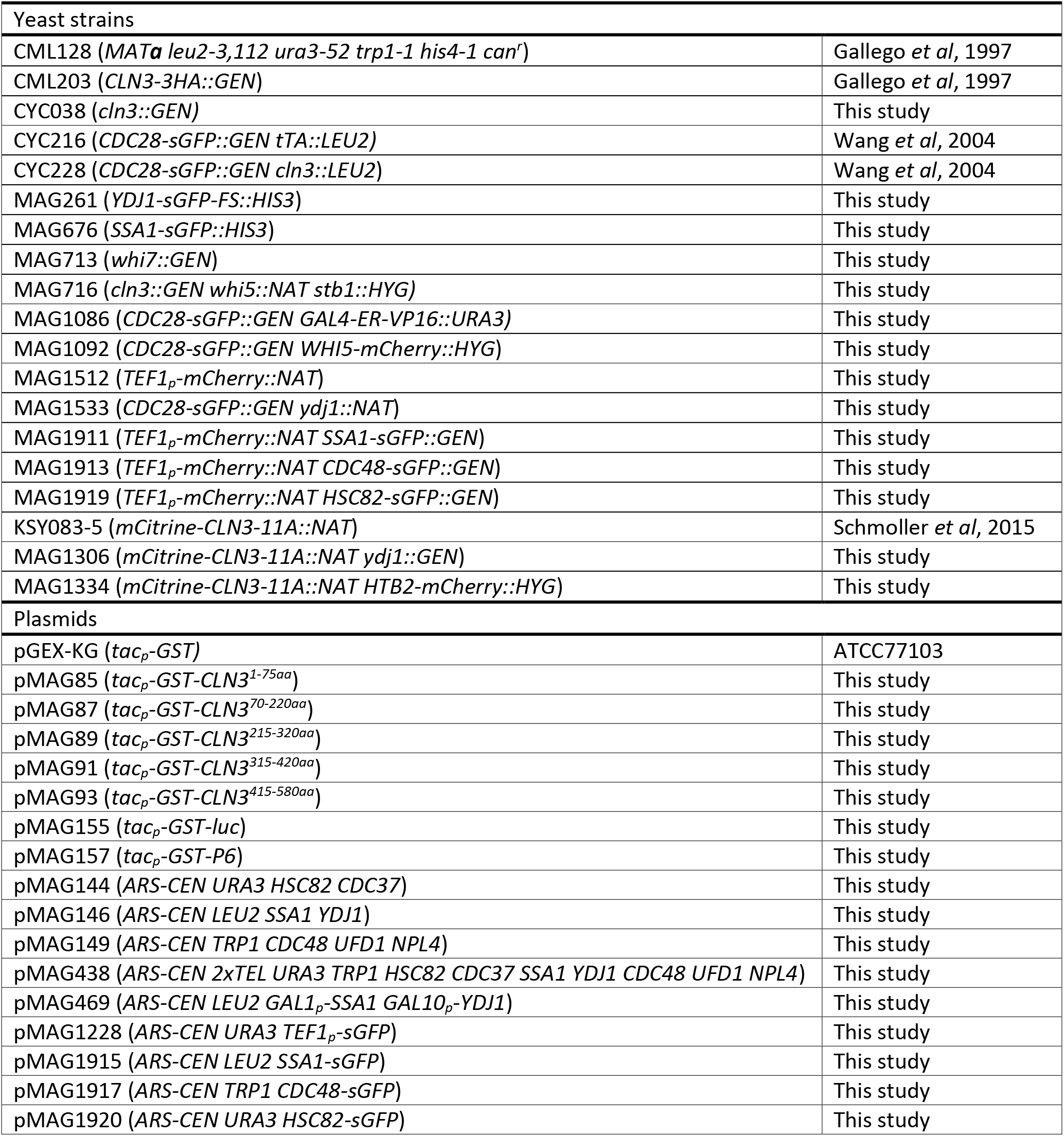
Yeast strains and plasmids.

